# *CAFE MOCHA:* An Integrated Platform for Discovering Clinically Relevant Molecular Changes in Cancer; an Example of Distant Metastasis and Recurrence-linked Classifiers in Head and Neck Squamous Cell Carcinoma

**DOI:** 10.1101/105577

**Authors:** Neeraja M Krishnan, I Mohanraj, Janani Hariharan, Binay Panda

## Abstract

**Background:** *CAFE MOCHA* (Clinical Association of Functionally Established MOlecular CHAnges) is an integrated GUI-driven computational and statistical framework to discover molecular signatures linked to a specific clinical attribute in a cancer type. We tested *CAFE MOCHA* in head and neck squamous cell carcinoma (HNSCC) for discovering a signature linked to distant metastasis and recurrence (MR) in 517 tumors from TCGA and validated the signature in 18 tumors from an independent cohort.

**Methods:** The platform integrates mutations and indels, gene expression, DNA methylation and copy number variations to discover a classifier first, predict an incoming tumour for the same by pulling defined class variables into a single framework that incorporates a coordinate geometry-based algorithm, called Complete Specificity Margin Based Clustering (CSMBC) with 100% specificity. *CAFE MOCHA* classifies an incoming tumour sample using either a matched normal or a built-in database of normal tissues. The application is packed and deployed using the *install4j* multi-platform installer.

**Results:** We tested *CAFE MOCHA* to discover a signature for distant metastasis and recurrence in HNSCC. The signature MR44 in HNSCC yielded 80% sensitivity and 100% specificity in the discovery stage and 100% sensitivity and 100% specificity in the validation stage.

**Conclusions:** *CAFE MOCHA* is a cancer type- and clinical attribute-agnostic computational and statistical framework to discover integrated molecular signature for a specific clinical attribute.

*CAFE MOCHA* is available in GitHub (https://github.com/binaypanda/CAFEMOCHA).

## Introduction

Cancer progression is linked to molecular changes at multiple levels, such as somatic mutations, gene expression, DNA methylation and copy number changes. In the last 5yrs, a large amount of data on key variants in multiple cancers has been generated by international consortia, like The Cancer Genome Atlas (TCGA) (Editorial, 2015; Weinstein *et al*, 2013), International Cancer Genome Consortium (ICGC) (Hudson *et al*, 2010) and individual laboratories, aiding our understanding of various cancers at molecular level. Utilizing the vast amount of molecular data from studies on tumour genomes, exomes, transcriptomes and methylomes, and linking those with data from genetic and functional studies will help find clinically relevant insights. Although there are existing databases and studies that combine molecular changes across cancer types (Deng *et al*, 2016; Huang *et al*, 2015; Netanely *et al*, 2016), studies linking the events explicitly within the same tumour type across a large number of samples to predict signatures for a specific clinical attribute are currently lacking.

Here, we describe a cancer type-agnostic computational and statistical framework with a user-friendly graphical-user-interface (GUI) to discover integrated signatures using six tumour-specific event types; somatic mutations and indels (mut), DNA copy number changes (cnv), gene expression changes (expr), 5-cytosine DNA methylation changes (meth), functional copy number changes (fcnv, where CNVs are linked to gene expression changes) and functional *cis*-regulatory DNA methylation changes (fmeth, where hyper- and hypo-methylation result in down- and up-regulation of effector gene expression respectively). *CAFE MOCHA*, in addition to classifying categorical events like mut and cnv, uses an algorithm called Complete Specificity Margin Based Clustering (CSMBC) to classify quantitative (expr and meth) and coupled events (fmeth and fcnv), and combines the event types using sample frequency and event priority filters, to produce integrated signatures describing the clinical variable with 100% specificity. We tested *CAFE MOCHA* to discover a signature linked with distant metastasis and recurrence in head and neck squamous cell carcinoma (HNSCC) using 434 tumours from TCGA as a discovery cohort, which was validated in an independent cohort of 18 tumours. The integrated signature MR44 for metastatic and recurrent tumours in HNSCC (MR44) yielded a sensitivity of 79.52% and a specificity of 100% in the discovery cohort and with 100% sensitivity and 100% specificity in the validation cohort.

## Materials and Methods

### Discovery module

*CAFE MOCHA* application workflow and the GUI are illustrated in Figure 1. *CAFE MOCHA* has two independent modules, discovery and prediction. In the discovery module, both somatic mutations (missense, nonsense and splice-site) and indels were considered under mut. A matrix of following values (thresholds) were considered for cnvs; -2: full copy deletion, -1: allelic deletion, 1: low-copy amplification, 2: high-copy amplification, and 0: lack of any copy number variation. For expression and methylation, RSEM (Li & Dewey, 2011) -normalized gene-wise intensity matrix and a probe-wise matrix of pre-processed β values (from Illumina 450k whole-genome methylation arrays), respectively, were used. The methylation data was pre-processed, including normalization and batch correction, as described previously (Krishnan *et al*, 2016). Tumour samples assayed for all the four events (mut, cnv, expr and meth) were considered for the discovery phase. The perturbed genes were passed through a functional filter (detailed below) before being used in the integrated analysis. In the case of methylation data, a sub-region filter was applied and only probes located in the gene promoters, transcription start site (TSS200 and TSS1500) and CpG islands were retained. Categorical events like mut and cnvs were linked to the chosen clinical attributes with complete specificity. Quantitative events such as expression (expr) and methylation (meth) were linked to clinical attributes using an algorithm called Complete Specificity Margin Based Clustering (CSMBC) algorithm (detailed below). A somatic filter based on an unpaired *t*-test (*P* < 0.05) was further used to retain only tumour-specific expr and meth events. Expression events were further coupled with meth and cnvs to generate coupled events (fmeth and fcnv) respectively, where meth and cnvs affected the target gene expression. We only considered hyper- and hypo-methylation events linked with the down- and up-regulation of downstream gene expression respectively.

**Figure 1:**
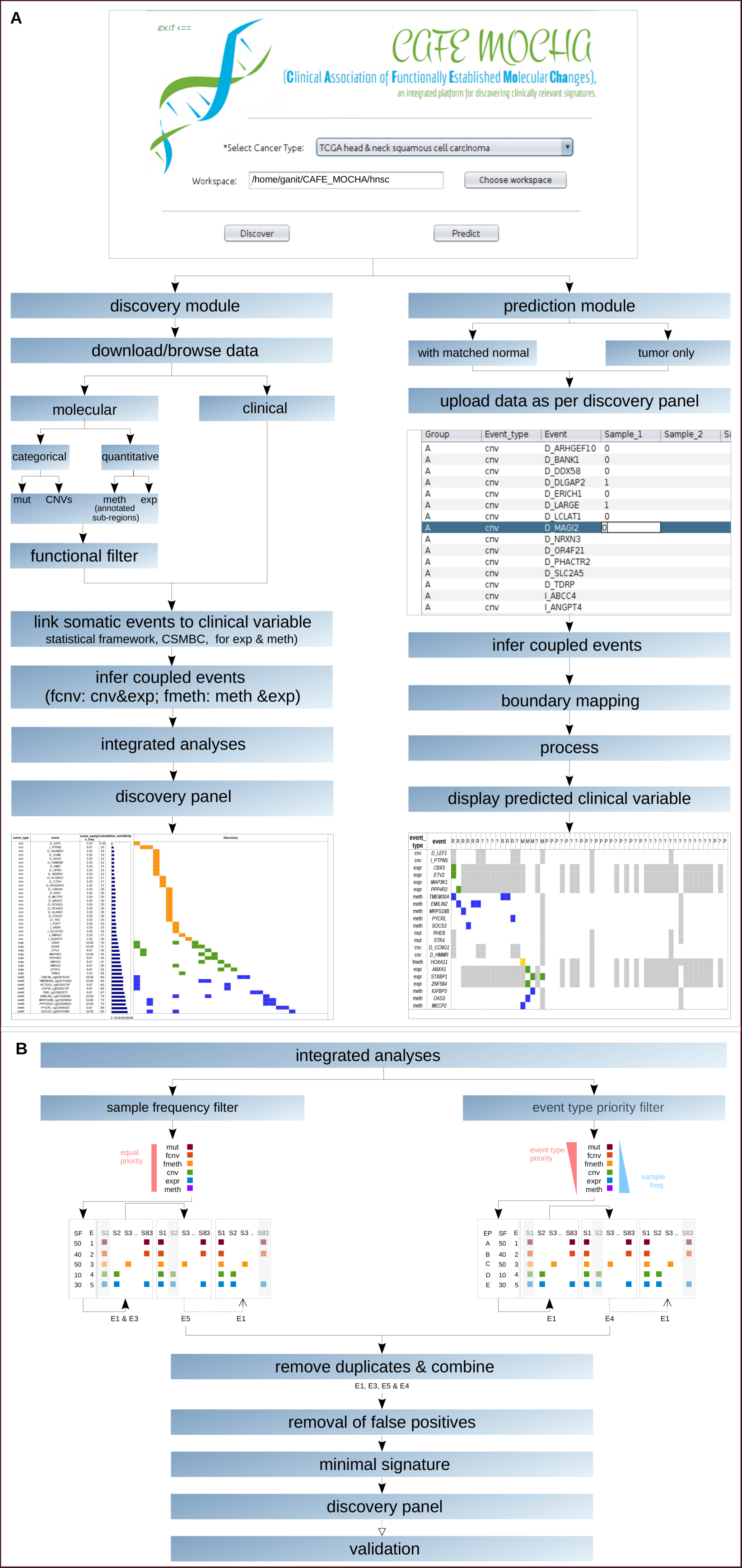
*CAFE MOCHA* application workflow and graphical-user-interface. A. Discovery, and Prediction modules. B. Integrated analyses.

### Data integration and making of the final signature

Using the ‘integrate’ feature, the individually discovered events were combined into an integrated signature with two different filters (Figure 1B). In the first, a tumour-specific sample frequency-based filter agnostic to any event type was applied. In the second, all event types’ priorities were determined by detection sensitivity first and then those events that yield highest priority followed by highest sample frequency were chosen. These two filters work contrarily, one retaining the high sample frequency quantitative and semi-quantitative event types (expr, meth, fmeth, fcnv) and another the lower-sample frequency categorical and semi-categorical event types (mut, cnv, fcnv). Selected events were considered for removal of false positives using samples lacking complete cross-platform overlap from TCGA and the independently provided datasets (used for confirmation in the second stage of the discovery stage). The number of confirmed events was further minimized using the sample frequency-based filter to constitute the discovery panel for that specific clinical attribute. This minimal signature is then subjected to an independent validation using tumours assayed for all the four events (mut, cnv, expr, meth) (Figure 1B). Finally, the discovered events were mapped to all of the sixteen cancer-related pathways from KEGG (Kanehisa *et al*, 2002; Kanehisa *et al*, 2004; Kanehisa *et al*, 2012) (hsa05200).

### CAFE MOCHA Interface

The interface for *CAFE MOCHA* was developed and deployed using *Netbeans IDE v 8.1* (https://netbeans.org/downloads/). The Graphical User Interface (GUI) was designed using JAVA AWT (http://www.javatpoint.com/java-awt) with SWING components (http://www.javatpoint.com/java-swing) with its native OS GUI and appropriate controls. The back end for this interface was built on a Linux-dependent platform with *R*, *BASH*, *BEDTools* (v2.3) and *PERL* as added dependencies. *CAFE MOCHA* requires installations of *R* package dependencies (minfi v1.18.2 (Aryee *et al*, 2014), wateRmelon v1.16.0 (Pidsley *et al*, 2013), IlluminaHumanMethylation450kmanifest v0.4.0, randomForest v4.6.12(Breiman, 2001), varSelRF v0.7.5, pheatmap v1.0.8, gplots v3.0.1, ggplot2 v2.1.0, reshape2 v1.4.1) to pre-process 450k *idat* files and generates the matrix of β values. The platform provides the user with an option to install the dependencies. Under the discovery module, mut, cnv, expr and meth data can be browsed and acquired locally.

### Functional filters

To ensure functional relevance of predicted tumour-specific molecular changes, we introduced various filters for different events to select genes of interest. We used IntOGen (Gundem *et al*, 2010) to select driver/potential driver mutations, MethHC (Huang *et al*, 2015) for known hyper- and hypomethylated genes in different cancer types, G2SBC (Mosca *et al*, 2010) for selecting important differentially expressed genes associated with breast cancer, The Human Protein Atlas (Uhlen *et al*, 2015) for functional proteins, the dbDEPC 2.0 database (He *et al*, 2012) for manually curated differentially expressed proteins in human cancer and miRTarBase (Chou *et al*, 2016; Hsu *et al*, 2011; Hsu *et al*, 2014) for experimentally validated miRNA-target gene associations. In the case of micro-RNA methylation, we considered those events where methylation of a microRNA correlated with a concomitant expression change in one or more of its target gene(s). In the case of fcnv, we considered where amplification and deletion in a gene was linked to its up- and down-regulation respectively in tumour sample compared to normal. Only full copy deletions (not allelic deletions) and amplifications were considered. Genes bearing categorical events in at least one sample that passed the functional filter were considered further.

### Complete Specificity Margin Based Clustering (CSMBC)

We devised a statistically framework, Complete Specificity Margin Based Clustering or CSMBC, for inferring the expression and the methylation boundaries of each cluster for a specific clinical attribute (Figure 2). In CSMBC, the number of clusters were fixed equivalent to the number of pre-defined clinical variables. For each cluster, axis-perpendicular boundaries passing through the most extreme outliers on the expression/methylation coordinate axes were determined first. The number of overlaps was dependent upon the number of boundaries. In the simplest case, there can be two sub-categories. For example, metastasis as a category has two sub-categories, metastatic (A); and non-metastatic (B). In this simplest case, there can only be one overlap 
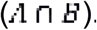
. Likewise, if a clinical attribute has three (A, B and C) sub-categories (for example recurrent, non-recurrent, can’t be defined) with four possible overlaps 
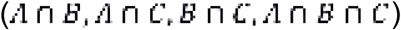
. Using the above logic defined for categories and sub-categories, all possible overlaps between clusters were identified first, and then the minimum and maximum values for various overlaps were estimated according to the following equations:

**Figure 2:**
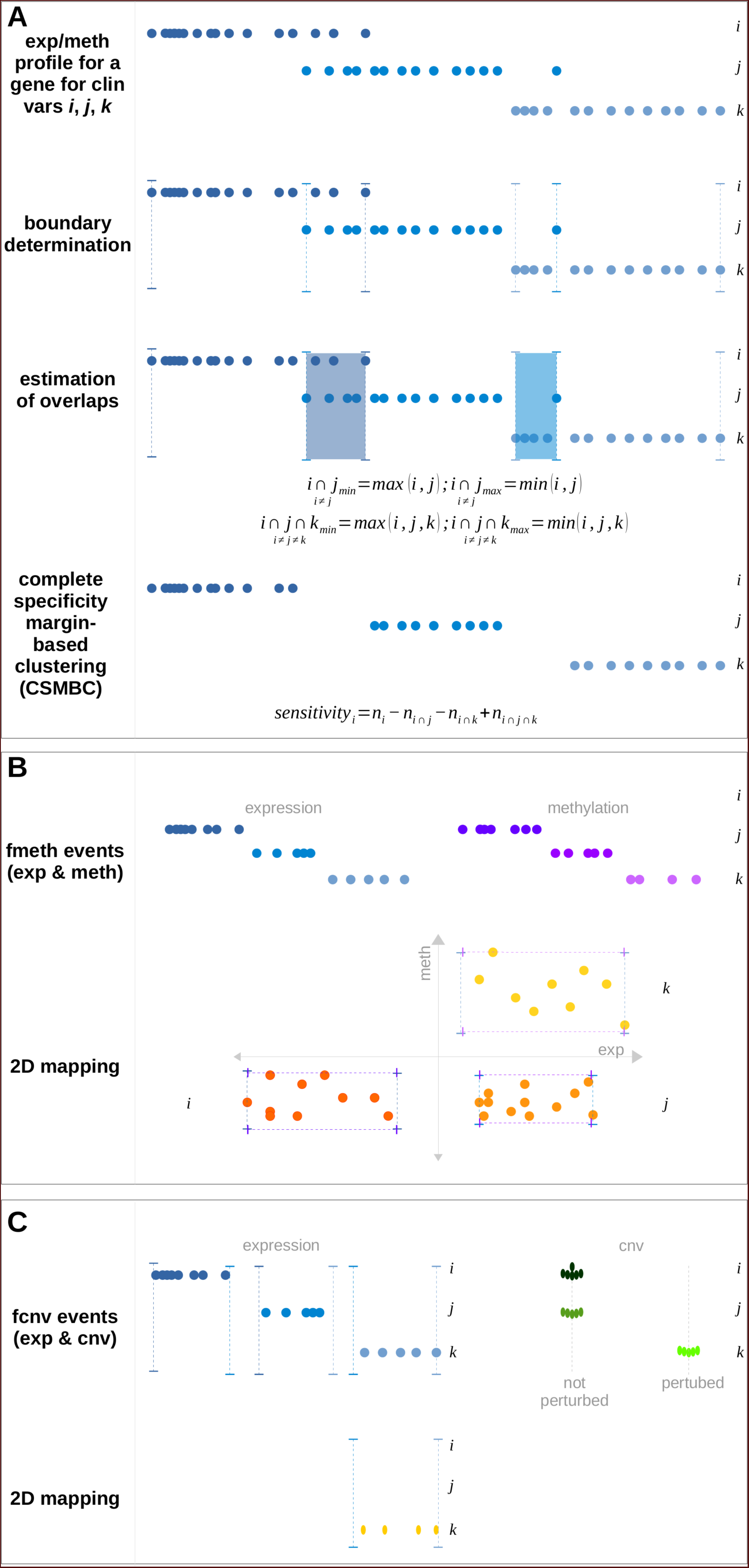
Complete specificity margin based clustering (CSMBC) algorithm for quantitative and coupled events. Clustering of quantitative and coupled events, for three clinical sub-categories *i*, *j* and *k* is demonstrated here. Panel A: linking of quantitative events such as expression or methylation to clinical sub-categories *i*, *j* and *k* (shades of blue). Boundaries are determined in a supervised manner, as the minimum and maximum limits of the quantitative ranges for each clinical sub-category. Overlaps are estimated as per the equations illustrated in the figure. Samples whose expression or methylation values lie outside the regions of overlap, 100% specific to a clinical sub-category are factored in towards the sensitivity of that gene for that clinical sub-category. Panel B: clinical linking of coupled events such as fmeth (shades of orange) resulting from a combination of two quantitative events (expr – shades of blue and meth – shades of purple). The expression event is mapped along the first dimension and the methylation event is mapped along the second, and the samples, which observe both, expression and methylation, boundaries, contribute to the sensitivity of that fmeth event for that clinical sub-category. Panel C: clinical linking of coupled events such as fcnv (shades of dark orange) resulting from a combination of one quantitative event (expression – shades of blue) and another categorical event (CNV – shades of green). Here, the fcnv event linked to a clinical sub- category is determined by both, expression boundaries for a clinical sub-group, and presence of CNVs for the same samples in that clinical sub-group.

For overlap between two sub-categories:

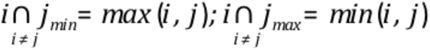

For overlap between three sub-categories:

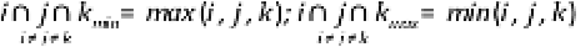

For overlap between four sub-categories:

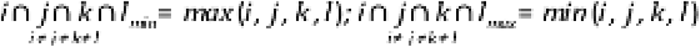

A sample was identified to be within 100% specificity margin of the cluster if it was not in any region of overlap with other clusters (Figure 2). The total number of all such samples, *i.e.*, sensitivity of that cluster, was estimated according to the equation below:

For two sub-categories:

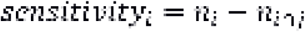

For three sub-categories:

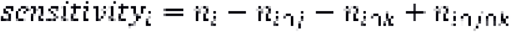

For four sub-categories:

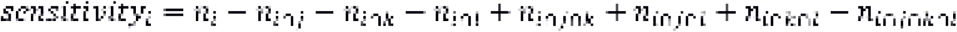

Finally, an unpaired *t*-test was performed between all tumour and all normal samples, and genes/probes with significantly different expr/meth values (*P* < 0.05) between the tumour and normal groups were retained.

### Prediction module

The prediction module has two options: one, where the user provides somatic mutations/indels, tumour-specific expr, meth and cnv information, and second, where the user has somatic mutations/indel/cnv data but does not have expr and meth data from the matched normal samples. In the later case, *CAFE MOCHA* uses a built in normal database (all normal samples from the TCGA for a particular cancer type) to compute expression and methylation values for genes in control samples from RNASeq for expression and whole genome arrays (Illumina 450k) for DNA methylation. For the prediction module, the data is entered as per the pre-defined format (same as discovery panel) specific to a cancer type and clinical attribute of interest (Figure 1). Based on the user input, coupled events (fmeth and fcnv) are determined and used for the prediction module.

### Testing CAFE MOCHA to discover metastatic/recurrent HNSCC signature

We tested *CAFE MOCHA* to discover integrated signature for metastatic and recurrent HNSCC tumours using tumours from TCGA dataset (n=434), followed by confirmation (removal of false positives, the second stage of the discovery phase) using two sample cohorts, 42 samples from TCGA and 37 samples from an independent cohort (Krishnan *et al*, 2015; Krishnan *et al*, 2016), where the information on at least one of the event type out of the 4 events (mut, cnv, expr and meth) for the same tumour was available (Table 1, Supplementary Tables S1 and S2). Finally, the discovery panel was validated in 18 samples from an independent sample cohort (Krishnan *et al*, 2015; Krishnan *et al*, 2016), where all the four events were assayed for all the tumours (Supplementary Table S3). For the discovery module, we downloaded the Broad automated somatic mutations and indel call file, copy number data with gistic2 threshold, Illumina HiSeqV2 RSEM (7) -normalized RNA-seq gene expression data, 450k DNA methylation data and the clinical data from the UCSC Xena Browser (http://tinyurl.com/jhmg9b9). The distant metastasis (M1) and recurrent tumours (RFS_IND = 1) were compared with non-metastatic (M0) and non-recurrent (RFS_IND = 0) tumours to derive a specific signature.

**Table 1.** Sample cohorts used in the discovery and validation modules of *CAFE MOCHA* for discovering metastatic/recurrent signature MR44 in head and neck squamous cell carcinoma (HNSCC).

| Discovery |  |  |  |
| --- | --- | --- | --- |
| Stage 1 | Source | TCGA HNSCC (http://tinyurl.com/jhmg9b9), data available on all the 4 event types (mut, CNV, exp, meth) on tumor:matched normal samples. |  |
|  | Total Number of tumors | 434 |  |
|  | Metastatic/Recurrent | 83 |  |
|  | Tumors without metastasis/recurrence | 351 |  |
| Stage 2 | Source | Data on at least one event type available from the same matched tumor:normal sample. |  |
|  |  | TCGA HNSCC | OTSCC (Krishnan *et al*, 2015; Krishnan *et al*, 2016) |
|  | Total number of tumors | 42 | 37 |
|  | Metastatic/Recurrent | 4 | 15 |
|  | Tumors without metastasis/recurrence | 38 | 22 |
|  | Adjacent normal tissue | 0 | 37 |
| Validation |  |  |  |
|  | Source | OTSCC (Krishnan *et al*, 2015; Krishnan *et al*, 2016) (matched tumor:normal data on all event types from the same tumors) |  |
|  | Total number of tumors | 18 |  |
|  | Metastatic/Recurrent | 3 |  |
|  | Tumors without metastasis/recurrence | 15 |  |
|  | Adjacent normal tissue | 18 |  |

## Results

### Discovery of distant metastasis/recurrence-associated molecular signature MR44 in HNSCC

*CAFE MOCHA* identified a total of 6649 events from individual event type discovery (8 somatic mutations, 1436 cnvs (506 amplifications and 982 full copy deletions), 1098 expression-related events, 4018 methylation events, 27 fcnv events (8 high copy amplifications and 19 full copy deletions) and 62 fmeth events), associated with metastasis and recurrence in HNSCC. Integration across discovered events was performed using two filters as described in the Methods section. The filters resulted in 79 (22 expression-related events and 57 methylated genes) and 171 (7 mut, 84 cnvs (46 amplifications and 38 full copy deletions), 24 expr, 2 meth, 25 fcnv events (8 high copy amplifications and 17 full copy deletions) and 29 fmeth events) events, individually, and 232 (7 mut, 84 cnvs (46 amplifications and 38 full copy deletions), 30 expr, 57 meth, 25 fcnv events (8 high copy amplifications and 17 full copy deletions) and 29 fmeth events), cumulatively after removing any redundancy (Supplementary Table S4).

Using data at the second stage of the discovery phase (confirmation stage), 98 false positives events were eliminated. The remaining 134 events (Supplementary Figure S1) (5 mut, 83 cnvs (46 amplifications and 37 full copy deletions), 8 expr, 9 meth, 24 fcnv events (8 high copy amplifications and 16 full copy deletions) and 5 fmeth events) were further minimized to a final discovery panel of 44 events (MR44) that included 2 mut, 15 cnvs (3 amplifications and 12 full copy deletions), 8 expr, 9 meth, 8 fcnv events (5 high copy amplifications and 3 full copy deletions) and 2 fmeth events) (Figure 3A). The signature was further validated using an independent cohort of 18 samples (3 M1/R and 15 non-MR) of an independent oral tongue squamous cell carcinoma (OTSCC) cohort, with 100% sensitivity and 100% specificity. 17 out of the 44 events in MR44 signature were validated in at least one sample in the validation cohort (Figure 3B).

**Figure 3:**
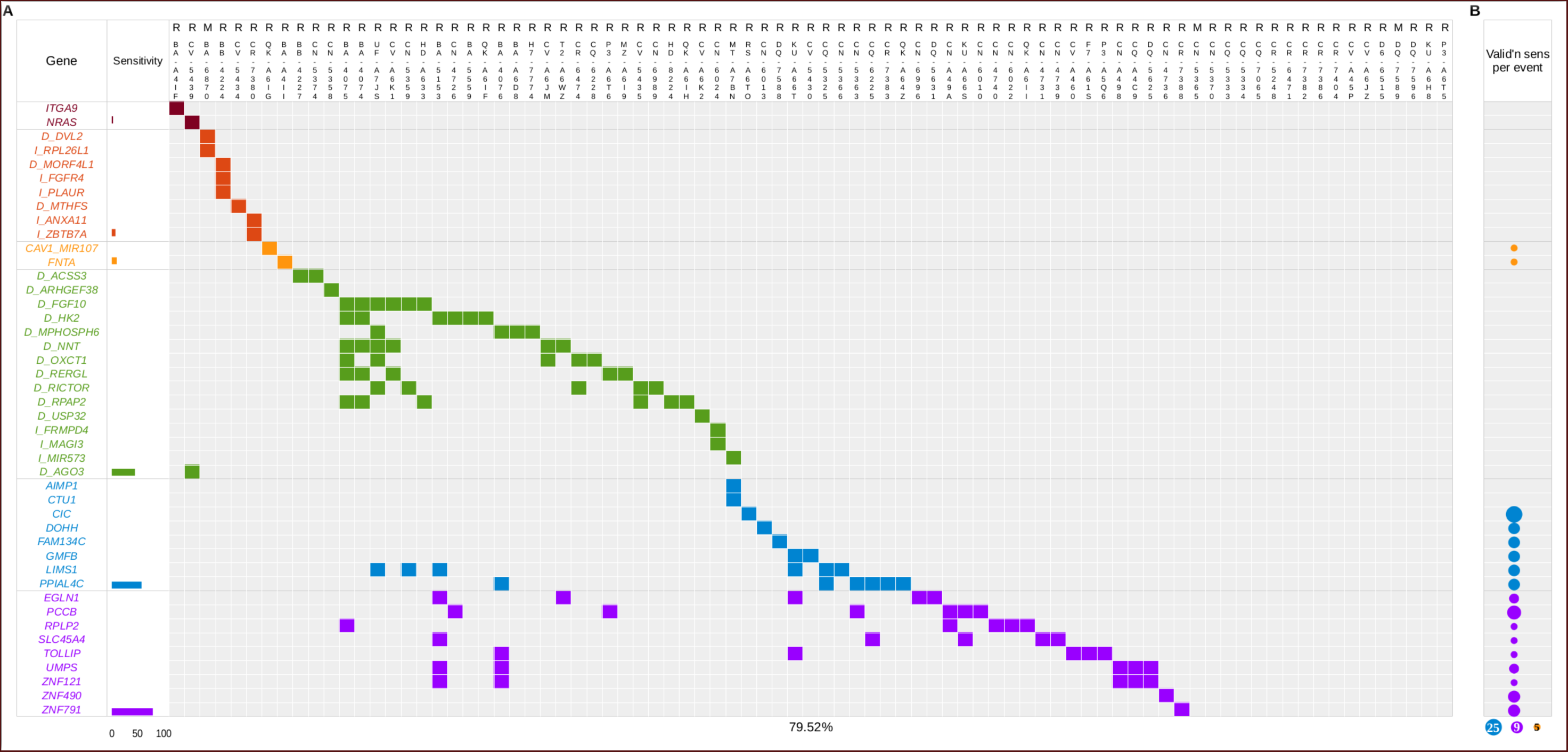
Discovery heatmap (A) and per-event validation sensitivity (B) of 44-gene integrated signature (MR44) for distant metastasis and recurrence in the HNSCC discovery set. An integrated signature associated with metastasis/recurrence was derived from combining six event types (mut: red, fmeth: dark orange, fcnv: orange, cnv: green, expr: blue and meth: purple). Cumulative sample frequency (%) for individual event types is represented as a histogram. The per-event validation sensitivity is represented as bubbles, where the bubble size is proportional to the sample frequency of that event in the validation cohort.

### Power of integration

We wanted to investigate the power of integration (where all the six different event types were used to derive the signature) over lesser order combinations or with individual event type, in terms of the detection sensitivity and the number of events required to achieve maximum specificity. As shown in Figure 4, the detection sensitivity showed a gradual enhancement when the number of event types increased from one to six. For single-event analyses, meth and both mut and fmeth provided with a maximum and minimum sensitivity respectively (37%, 7%, Figure 4A). The number of CNVs required to achieve the highest possible level of sensitivity in a single-event analysis was the highest (83 events with 33% sensitivity). When using more than one event type, we obtained an increase in sensitivity to various levels gradually from a single-event to a six-events signature (Figure 4). The number of CNVs and both mut and fmeth required to get similar level of sensitivity was highest and lowest respectively for all combinations. Highest sensitivity (80%) of detection was observed when all the six different event types were used to produce the integrated signature (Figure 4F). The impact of event integration on detection sensitivity was observed to be greatest when the integration was performed in the following order: meth + cnvs + expr + fmeth + fcnvs + mut. Each incremental integration in this order caused a gradual increase in sensitivity (37.35 → 56.63 → 68.67 → 74.70 → 77.11 → 79.52, Figure 4). The actual events for all combinations and individual event type analyses are illustrated in Supplementary Figure S1.

**Figure 4:**
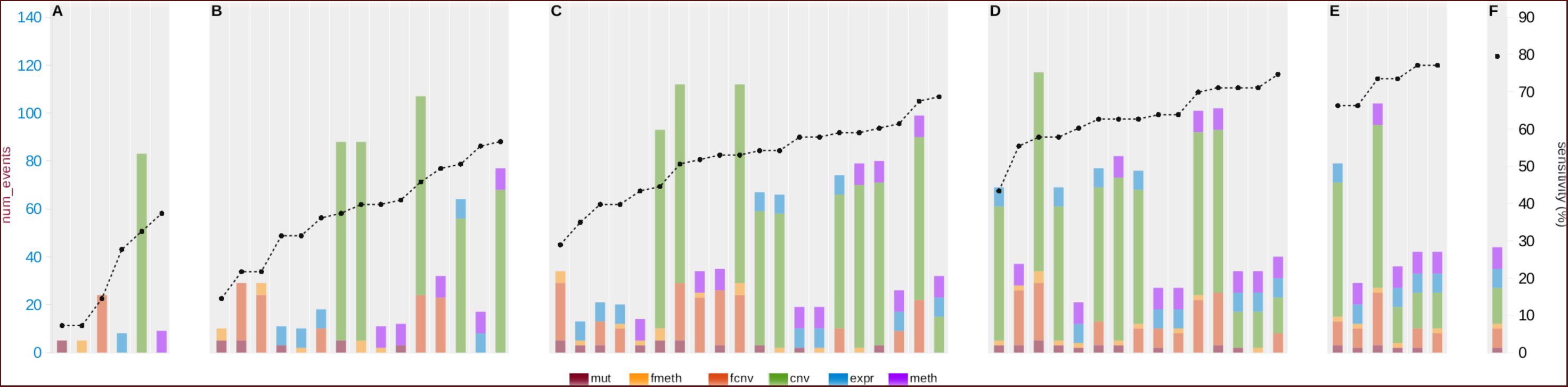
Comparison of detection sensitivities and total numbers of events required achieving the sensitivity in one (A), two (B), three (C), four (D), five (E) and all six (F) event types. Utilizing all the six events (F) shows the power of integration both on sensitivity of detection (black dot) and the number of events (colored bars for all the six individual event types) required to attain the sensitivity.

## Discussion

Integrating multiple molecular events is a strategy proposed to identify master regulators in cancer (Gevaert & Plevritis, 2013; Thingholm *et al*, 2016). However, a meaningful integration should be incremental, with each level filtering out the unnecessary and only retaining the functionally relevant associations. Furthermore, integration of data should link a specific signature to clinically relevant tumour attributes in order for making meaningful conclusions. Integrated analyses often fail to establish the link between cancer-associated changes within the same tumour types across cohorts, perhaps due to the low overlapping of clinical characteristics across cohorts and low reproducibility of results across laboratories, geographies and discovery platforms. Additionally, data on multiple events; genetic, transcriptomic and epigenetic changes, from the same sets of tumour:matched normal samples are often not available, making it difficult to discover truly integrated signatures. Nevertheless, such a discovery set, where all the events are assayed for all tumour:normal pairs within the same cohort, is functionally more meaningful. As multidimensionality increases with the availability of more data, especially from the large consortia like TCGA and ICGC, it will become imperative to design and implement easy-to-use and robust web-based platforms to discover (using a pre-defined training set) multi-gene and multi-platform classifiers for a particular clinical attribute and predict the same for an incoming new tumour sample. Currently, such a platform is lacking that can discover tumour-specific genome-wide functional and somatic molecular changes linked to a specific clinical attribute, using a sizeable multi-dimensional dataset. Keeping this in mind, we devised *CAFE MOCHA*, an automated and integrated framework to discover meaningful, functional and somatic molecular changes in a cancer type that links the signature(s) to a specific clinical attribute. *CAFE MOCHA* is designed to use both user-defined/generated and publicly available tumour and matched normal data.

Additionally, the prediction module of *CAFE MOCHA* uses a backend database of unmatched normal samples to increase the prediction scope of the tool for tumours where matched normal samples are not available, especially where expression and methylation data is not available from matched normal samples. *CAFE MOCHA* is a cancer type- and clinical attribute-agnostic tool and has the ability to pull data on several event types under a single framework. *CAFE MOCHA* classifies events into three different types, categorical (mutations and CNV), quantitative (expression and methylation) and coupled/linked (functional CNVs and functional methylation) and uses an algorithm called complete specificity margin based clustering (CSMBC), a fully supervised approach, to identify clinically linked quantitative and coupled events. CSMBC is a modification of the Large Rectangle Margin Learning (LRML) approach described previously (kirmse & Petersohn, 2011) and is conceptually similar to fast boxes which take the ‘characterize’ and then ‘discriminate’ approach of classification (Goh & Rubin, 2014). The input for CSMBC is a continuous function spread across a single dimension, either expression or methylation. For both these variables, an unsupervised clustering, which requires *a priori* knowledge of the quantitative expression or methylation map, is not feasible since these are continuous variables. The selection of boundary margins, according to the chosen clinical variables on the other hand does not require any prior knowledge of the quantitative spread of these events. Additionally, supervised clustering is faster than unsupervised clustering since it does not require iterative computing and does not require re-estimation of the cluster mean. Nevertheless, as pointed out in the past (Richards, 2013), a hybrid semi-supervised approach that uses the results of an unsupervised approach such as the *k*-means or Expectation Maximization (EM) as training areas for a supervised classification, might combine the advantages of both supervised and unsupervised classification approaches. A weighted sensitivity method (Gao, 2007; Iwamoto & Pusztai, 2010) that weighs the classification sensitivity of a sample based on its proximity to the nearest boundary, would further add confidence perhaps at the cost of sensitivity, particularly while predicting the clinical status of an incoming sample. While we plan to develop *CAFE MOCHA* taking these modifications into account in the future, the current implementation benefits from ensuring complete specificity for each clinical attribute-associated molecular event. One of the limitations of CSMBC is that its specificity is entirely dependent on the denominator, *i.e.,* sample numbers and therefore, may be sub-optimal in a scenario where fewer numbers of tumour samples are present in the discovery set.

HNSCC are the sixth leading cause of cancer worldwide with 5-year survival of less than 50% (Ferlay *et al*, 2010; Mishra & Meherotra, 2014). Recent high-throughput studies employing computational methods have identified various genetic, transcriptomic and epigenetic changes from different subsites of HNSCC from different geographies (Agrawal *et al*, 2011; Cancer Genome Atlas, 2015; India Project Team of the International Cancer Genome, 2013; Krishnan *et al*, 2015; Pickering *et al*, 2013). Early-stage patients with HNSCC are usually treated using a single modality like surgery or radiotherapy whereas advanced-stage patients benefit from multi-modality therapies (Ridge *et al*, 2016). In head and neck tumours, identifying patients with tumours *a priori* that may undergo distant metastasis or loco-regional recurrence using primary-tumour-derived molecular signature will help manage patients better. *CAFE MOCHA* was tested using a robust dataset of 434 HNSCC tumours from TCGA where data on all four molecular events were available. Despite the MR44 being an integrated signature, drawn from 83 metastatic/recurrent tumours with four different somatic events (mut, cnv, expr and meth) and two coupled events (fmeth and fcnv) available on all the tumours, the discovery sensitivity did not attain close to 100% (sensitivity was 79.52%). This means that ~20% of the metastatic/recurrent tumours could not be classified using the integrated MR44 signature. This could indicate two possibilities: first, there are other types of tumour-specific changes that need to be assayed and accounted for in order to derive a truly complete distant metastasis and recurrence-associated signature, and/or, second: the sample size of 83 metastatic/recurrent tumours was not sufficient (the predictive power of discovery was not optimum). The fact that the validation sensitivity of MR44 was 100% in an independent cohort does not deter from either of these conclusions as the number of tumours in the validation set was small (*n* = 18).

About half of the genes in the MR44 signature have been reported previously to be associated with prognosis, survival, recurrence and distant metastasis for various cancer types. For example, *ITGA9* is reported as a tumour suppressor gene in non-small cell lung cancer (Pastuszak-Lewandoska *et al*, 2016) and is involved in cell migration and invasion in melanoma (Zhang *et al*, 2016). Additionally, epigenetic inactivation of *ITGA9* is linked with its expression in nasopharyngeal and breast cancer (Mostovich *et al*, 2011; Nawaz *et al*, 2015). *NRAS* is linked with survival of patients with liver metastases in colorectal cancer (Vauthey *et al*, 2013) and acts as a prognostic factor in metastatic melanoma (Jakob *et al*, 2012). The chromosomal region containing *DVL2* gene frequently undergoes LOH in patients with colorectal tumours (Kurashina *et al*, 2008). *RPL26A1* expression was a prognostic marker in ER-positive breast tumours (Wang & Zhang, 2007) and liver metastases in colorectal cancer (Nakamura & Furukawa, 2003). *MORF4L1* was found to be one of the candidate genes with copy number reductions in breast cancer (Chen *et al*, 2007). *FGFR4* profile was observed to be prognostically relevant in squamous cell carcinoma of the mouth and oropharynx (Dutra *et al*, 2012), esophagus (Shim *et al*, 2016), and gastric cancer (Murase *et al*, 2014). *ANXA11* expression is one of the prognostic markers in prostate cancer (Chandran *et al*, 2007; Tsai *et al*, 2013). *ZBTB7A* plays a role in suppressing tumour metastasis in gastric cancer (Shi *et al*, 2015). Loss of stromal *CAV1* expression predicts poor survival in colorectal cancer (Zhao *et al*, 2015). *ARHGEF38* expression significantly differed between high and low recurrence-free survival groups in prostrate cancer (Tandefelt *et al*, 2013). *OXCT1* was shown to act as an oncogene in human breast cancer cells (Martinez-Outschoorn *et al*, 2012). *RICTOR* regulates cancer cell metastases in breast cancer cell lines (Zhang *et al*, 2010). *RPAP2* associates with breast cancer recurrence (Baker *et al*, 2014). mIR573 inhibits prostate cancer metastases (Wang *et al*, 2015). *AIMP1* is down regulated in gastric and colorectal cancer (Kim *et al*, 2014). *FAM134C* is a cancer-relapse marker in breast and ovarian cancer (Guo, 2011). *LIMS1* (PINCH signalling) has been linked to distant metastasis in pancreatic stromal cells (Scaife *et al*, 2010). *EGLN1* (commonly referred to as *PHD2*) is an oxygen sensor that promotes breast cancer metastases (Kuchnio *et al*, 2015). *RPLP2* is one of the 10-gene prognostic markers for gastric cancer (Zhang *et al*, 2011). *TOLLIP* was found to be significantly and differentially expressed in colorectal metastatic cells (Barderas *et al*, 2013). *ZNF490* was found to be associated with breast cancer recurrence (Baker *et al*, 2013). MR44 contains genes implicated in MTOR, MAPK and PI3K-AKT signalling pathways via *DVL2*, *FGF10*, *FGFR4*, *RICTOR*, *ITGA9*, *NRAS* and *CAV1* genes (Supplementary Table S4).

Although the discovery module of *CAFE MOCHA* uses 450k methylation array and RSEM-normalized RNASeq expression data, the predict module is not restricted to the assays being performed on the same platform so long as the user provides data in the same format for a matched normal. Future versions of *CAFE MOCHA* will incorporate cross-platform converter wherein the user can deposit data generated with multiple platforms and in multiple formats, thus making the tool truly platform agnostic. Additionally, *CAFE MOCHA* has the ability to use an underlying tissue-specific normal database, especially where expression and methylation data is not available, for un-matched normal samples. However, if data for a matched normal is not available, the user must use the same assays and formats, used in the discovery module to minimize assay/platform related errors.

## Supplementary Data Legend

**Supplementary Table S1:** TCGA HNSCC tumor samples (*n* = 434) and clinical attributes used for the discovery of MR44.

| SampleID | R (Loco-regional recurrence); M1(metastasis); M0 (Non-metastatic/Non-recurrent) tumors; N (Solid Tissue Normals). |
| --- | --- |
| TCGA-BA-4074-01 | R |
| TCGA-BA-4075-01 | R |
| TCGA-BA-4076-01 | R |
| TCGA-BA-4077-01 | M0 |
| TCGA-BA-4078-01 | M0 |
| TCGA-BA-5149-01 | M0 |
| TCGA-BA-5151-01 | M0 |
| TCGA-BA-5152-01 | M0 |
| TCGA-BA-5153-01 | R |
| TCGA-BA-5555-01 | M0 |
| TCGA-BA-5556-01 | M0 |
| TCGA-BA-5557-01 | M0 |
| TCGA-BA-5558-01 | M0 |
| TCGA-BA-5559-01 | R |
| TCGA-BA-6868-01 | M0 |
| TCGA-BA-6869-01 | M0 |
| TCGA-BA-6870-01 | M1 |
| TCGA-BA-6871-01 | M0 |
| TCGA-BA-6872-01 | M0 |
| TCGA-BA-6873-01 | M0 |
| TCGA-BA-7269-01 | M0 |
| TCGA-BA-A4IF-01 | R |
| TCGA-BA-A4IH-01 | M0 |
| TCGA-BA-A4II-01 | R |
| TCGA-BA-A6D8-01 | R |
| TCGA-BA-A6DA-01 | M0 |
| TCGA-BA-A6DB-01 | M0 |
| TCGA-BA-A6DD-01 | M0 |
| TCGA-BA-A6DE-01 | M0 |
| TCGA-BA-A6DI-01 | M0 |
| TCGA-BA-A6DJ-01 | M0 |
| TCGA-BA-A6DL-01 | M0 |
| TCGA-BB-4217-01 | M0 |
| TCGA-BB-4223-01 | M0 |
| TCGA-BB-4224-01 | R |
| TCGA-BB-4225-01 | M0 |
| TCGA-BB-4227-01 | R |
| TCGA-BB-4228-01 | M0 |
| TCGA-BB-7861-01 | M0 |
| TCGA-BB-7862-01 | M0 |
| TCGA-BB-7863-01 | M0 |
| TCGA-BB-7864-01 | M0 |
| TCGA-BB-7870-01 | M0 |
| TCGA-BB-7871-01 | M0 |
| TCGA-BB-7872-01 | M0 |
| TCGA-C9-A47Z-01 | M0 |
| TCGA-C9-A480-01 | M0 |
| TCGA-CN-4723-01 | M0 |
| TCGA-CN-4725-01 | M0 |
| TCGA-CN-4726-01 | R |
| TCGA-CN-4727-01 | M0 |
| TCGA-CN-4728-01 | M0 |
| TCGA-CN-4729-01 | M0 |
| TCGA-CN-4730-01 | M0 |
| TCGA-CN-4731-01 | R |
| TCGA-CN-4733-01 | M0 |
| TCGA-CN-4735-01 | M0 |
| TCGA-CN-4736-01 | R |
| TCGA-CN-4737-01 | M0 |
| TCGA-CN-4738-01 | M0 |
| TCGA-CN-4739-01 | R |
| TCGA-CN-4740-01 | R |
| TCGA-CN-4741-01 | M0 |
| TCGA-CN-4742-01 | M0 |
| TCGA-CN-5355-01 | M0 |
| TCGA-CN-5356-01 | M0 |
| TCGA-CN-5358-01 | R |
| TCGA-CN-5359-01 | R |
| TCGA-CN-5360-01 | M0 |
| TCGA-CN-5363-01 | R |
| TCGA-CN-5364-01 | M0 |
| TCGA-CN-5365-01 | M1 |
| TCGA-CN-5366-01 | R |
| TCGA-CN-5367-01 | M0 |
| TCGA-CN-5369-01 | M0 |
| TCGA-CN-5370-01 | R |
| TCGA-CN-5373-01 | M0 |
| TCGA-CN-5374-01 | R |
| TCGA-CN-6010-01 | R |
| TCGA-CN-6011-01 | M0 |
| TCGA-CN-6012-01 | M0 |
| TCGA-CN-6013-01 | R |
| TCGA-CN-6016-01 | M0 |
| TCGA-CN-6017-01 | M0 |
| TCGA-CN-6018-01 | M0 |
| TCGA-CN-6019-01 | M0 |
| TCGA-CN-6020-01 | M0 |
| TCGA-CN-6021-01 | M0 |
| TCGA-CN-6022-01 | R |
| TCGA-CN-6023-01 | M0 |
| TCGA-CN-6024-01 | R |
| TCGA-CN-6988-01 | M0 |
| TCGA-CN-6989-01 | R |
| TCGA-CN-6992-01 | M0 |
| TCGA-CN-6994-01 | M0 |
| TCGA-CN-6995-01 | M0 |
| TCGA-CN-6996-01 | R |
| TCGA-CN-6997-01 | M0 |
| TCGA-CN-6998-01 | M0 |
| TCGA-CN-A498-01 | R |
| TCGA-CN-A49A-01 | R |
| TCGA-CN-A642-01 | M0 |
| TCGA-CN-A6V3-01 | M0 |
| TCGA-CQ-5323-01 | M0 |
| TCGA-CQ-5324-01 | M0 |
| TCGA-CQ-5325-01 | R |
| TCGA-CQ-5326-01 | M0 |
| TCGA-CQ-5327-01 | M0 |
| TCGA-CQ-5329-01 | M0 |
| TCGA-CQ-5330-01 | M0 |
| TCGA-CQ-5331-01 | M0 |
| TCGA-CQ-5332-01 | M0 |
| TCGA-CQ-5333-01 | R |
| TCGA-CQ-5334-01 | R |
| TCGA-CQ-6218-01 | M0 |
| TCGA-CQ-6220-01 | M0 |
| TCGA-CQ-6221-01 | M0 |
| TCGA-CQ-6223-01 | M0 |
| TCGA-CQ-6224-01 | M0 |
| TCGA-CQ-6225-01 | R |
| TCGA-CQ-6227-01 | M0 |
| TCGA-CQ-6228-01 | R |
| TCGA-CQ-6229-01 | M0 |
| TCGA-CQ-7063-01 | M0 |
| TCGA-CQ-7065-01 | R |
| TCGA-CQ-7067-01 | M0 |
| TCGA-CQ-7068-01 | M0 |
| TCGA-CQ-7069-01 | M0 |
| TCGA-CQ-7071-01 | M0 |
| TCGA-CQ-7072-01 | M0 |
| TCGA-CQ-A4C6-01 | M0 |
| TCGA-CQ-A4C7-01 | M0 |
| TCGA-CQ-A4C9-01 | R |
| TCGA-CQ-A4CB-01 | M0 |
| TCGA-CQ-A4CD-01 | M0 |
| TCGA-CQ-A4CE-01 | M0 |
| TCGA-CQ-A4CG-01 | M0 |
| TCGA-CQ-A4CH-01 | M0 |
| TCGA-CR-5243-01 | M0 |
| TCGA-CR-5247-01 | M0 |
| TCGA-CR-5248-01 | R |
| TCGA-CR-5249-01 | M0 |
| TCGA-CR-6467-01 | M0 |
| TCGA-CR-6470-01 | M0 |
| TCGA-CR-6471-01 | R |
| TCGA-CR-6472-01 | M0 |
| TCGA-CR-6473-01 | M0 |
| TCGA-CR-6474-01 | R |
| TCGA-CR-6477-01 | M0 |
| TCGA-CR-6478-01 | M0 |
| TCGA-CR-6480-01 | M0 |
| TCGA-CR-6481-01 | M0 |
| TCGA-CR-6482-01 | M0 |
| TCGA-CR-6484-01 | M0 |
| TCGA-CR-6487-01 | M0 |
| TCGA-CR-6488-01 | M0 |
| TCGA-CR-6491-01 | M0 |
| TCGA-CR-6492-01 | M0 |
| TCGA-CR-6493-01 | M0 |
| TCGA-CR-7364-01 | M0 |
| TCGA-CR-7365-01 | M0 |
| TCGA-CR-7367-01 | M0 |
| TCGA-CR-7368-01 | M0 |
| TCGA-CR-7369-01 | M0 |
| TCGA-CR-7370-01 | M0 |
| TCGA-CR-7371-01 | M0 |
| TCGA-CR-7372-01 | M0 |
| TCGA-CR-7373-01 | M0 |
| TCGA-CR-7374-01 | M0 |
| TCGA-CR-7376-01 | M0 |
| TCGA-CR-7377-01 | M0 |
| TCGA-CR-7379-01 | M0 |
| TCGA-CR-7380-01 | R |
| TCGA-CR-7382-01 | R |
| TCGA-CR-7383-01 | R |
| TCGA-CR-7385-01 | M0 |
| TCGA-CR-7386-01 | R |
| TCGA-CR-7388-01 | R |
| TCGA-CR-7389-01 | M0 |
| TCGA-CR-7390-01 | M0 |
| TCGA-CR-7391-01 | M0 |
| TCGA-CR-7392-01 | M0 |
| TCGA-CR-7393-01 | M0 |
| TCGA-CR-7394-01 | M0 |
| TCGA-CR-7395-01 | M0 |
| TCGA-CR-7397-01 | M0 |
| TCGA-CR-7398-01 | M0 |
| TCGA-CR-7399-01 | M0 |
| TCGA-CR-7401-01 | M0 |
| TCGA-CR-7402-01 | M0 |
| TCGA-CR-7404-01 | R |
| TCGA-CV-5430-01 | R |
| TCGA-CV-5432-01 | M0 |
| TCGA-CV-5434-01 | R |
| TCGA-CV-5435-01 | R |
| TCGA-CV-5436-01 | M0 |
| TCGA-CV-5439-01 | R |
| TCGA-CV-5440-01 | M0 |
| TCGA-CV-5441-01 | M0 |
| TCGA-CV-5442-01 | M0 |
| TCGA-CV-5443-01 | M0 |
| TCGA-CV-5444-01 | M0 |
| TCGA-CV-5966-01 | M0 |
| TCGA-CV-5970-01 | M0 |
| TCGA-CV-5971-01 | M0 |
| TCGA-CV-5973-01 | M0 |
| TCGA-CV-5976-01 | M0 |
| TCGA-CV-5977-01 | M0 |
| TCGA-CV-5978-01 | M0 |
| TCGA-CV-5979-01 | M0 |
| TCGA-CV-6003-01 | M0 |
| TCGA-CV-6433-01 | M0 |
| TCGA-CV-6436-01 | M0 |
| TCGA-CV-6441-01 | M0 |
| TCGA-CV-6933-01 | M0 |
| TCGA-CV-6934-01 | M0 |
| TCGA-CV-6935-01 | M0 |
| TCGA-CV-6936-01 | M0 |
| TCGA-CV-6937-01 | M0 |
| TCGA-CV-6938-01 | M0 |
| TCGA-CV-6939-01 | M0 |
| TCGA-CV-6940-01 | M0 |
| TCGA-CV-6941-01 | M0 |
| TCGA-CV-6942-01 | M0 |
| TCGA-CV-6943-01 | M0 |
| TCGA-CV-6945-01 | M0 |
| TCGA-CV-6948-01 | M0 |
| TCGA-CV-6950-01 | M0 |
| TCGA-CV-6951-01 | M0 |
| TCGA-CV-6952-01 | M0 |
| TCGA-CV-6953-01 | M0 |
| TCGA-CV-6954-01 | M0 |
| TCGA-CV-6955-01 | M0 |
| TCGA-CV-6956-01 | M0 |
| TCGA-CV-6959-01 | M0 |
| TCGA-CV-6960-01 | M0 |
| TCGA-CV-6961-01 | M0 |
| TCGA-CV-6962-01 | M0 |
| TCGA-CV-7089-01 | M0 |
| TCGA-CV-7090-01 | M0 |
| TCGA-CV-7091-01 | M0 |
| TCGA-CV-7095-01 | M0 |
| TCGA-CV-7097-01 | M0 |
| TCGA-CV-7099-01 | M0 |
| TCGA-CV-7100-01 | M0 |
| TCGA-CV-7101-01 | M0 |
| TCGA-CV-7102-01 | M0 |
| TCGA-CV-7103-01 | M0 |
| TCGA-CV-7104-01 | M0 |
| TCGA-CV-7177-01 | M0 |
| TCGA-CV-7178-01 | M0 |
| TCGA-CV-7180-01 | M0 |
| TCGA-CV-7183-01 | M0 |
| TCGA-CV-7235-01 | M0 |
| TCGA-CV-7236-01 | M0 |
| TCGA-CV-7238-01 | M0 |
| TCGA-CV-7242-01 | M0 |
| TCGA-CV-7243-01 | M0 |
| TCGA-CV-7245-01 | M0 |
| TCGA-CV-7247-01 | M0 |
| TCGA-CV-7248-01 | M0 |
| TCGA-CV-7250-01 | M0 |
| TCGA-CV-7252-01 | M0 |
| TCGA-CV-7253-01 | M0 |
| TCGA-CV-7254-01 | M0 |
| TCGA-CV-7255-01 | M0 |
| TCGA-CV-7261-01 | M0 |
| TCGA-CV-7263-01 | M0 |
| TCGA-CV-7406-01 | M0 |
| TCGA-CV-7407-01 | M0 |
| TCGA-CV-7409-01 | M0 |
| TCGA-CV-7410-01 | M0 |
| TCGA-CV-7411-01 | M0 |
| TCGA-CV-7413-01 | M0 |
| TCGA-CV-7414-01 | M0 |
| TCGA-CV-7415-01 | M0 |
| TCGA-CV-7416-01 | M0 |
| TCGA-CV-7418-01 | M0 |
| TCGA-CV-7421-01 | M0 |
| TCGA-CV-7422-01 | M0 |
| TCGA-CV-7423-01 | M0 |
| TCGA-CV-7424-01 | M0 |
| TCGA-CV-7425-01 | M0 |
| TCGA-CV-7427-01 | M0 |
| TCGA-CV-7429-01 | M0 |
| TCGA-CV-7430-01 | M0 |
| TCGA-CV-7432-01 | M0 |
| TCGA-CV-7433-01 | M0 |
| TCGA-CV-7434-01 | M0 |
| TCGA-CV-7435-01 | M0 |
| TCGA-CV-7437-01 | M0 |
| TCGA-CV-7438-01 | M0 |
| TCGA-CV-7440-01 | M0 |
| TCGA-CV-A45O-01 | M0 |
| TCGA-CV-A45P-01 | R |
| TCGA-CV-A45Q-01 | M0 |
| TCGA-CV-A45R-01 | M0 |
| TCGA-CV-A45T-01 | M0 |
| TCGA-CV-A45U-01 | M0 |
| TCGA-CV-A45V-01 | M0 |
| TCGA-CV-A45W-01 | M0 |
| TCGA-CV-A45X-01 | M0 |
| TCGA-CV-A45Y-01 | M0 |
| TCGA-CV-A45Z-01 | M0 |
| TCGA-CV-A460-01 | R |
| TCGA-CV-A461-01 | M0 |
| TCGA-CV-A463-01 | M0 |
| TCGA-CV-A464-01 | M0 |
| TCGA-CV-A465-01 | M0 |
| TCGA-CV-A468-01 | M0 |
| TCGA-CV-A6JD-01 | M0 |
| TCGA-CV-A6JE-01 | M0 |
| TCGA-CV-A6JM-01 | R |
| TCGA-CV-A6JN-01 | M0 |
| TCGA-CV-A6JO-01 | M0 |
| TCGA-CV-A6JT-01 | M0 |
| TCGA-CV-A6JU-01 | M0 |
| TCGA-CV-A6JY-01 | M0 |
| TCGA-CV-A6JZ-01 | R |
| TCGA-CV-A6K0-01 | M0 |
| TCGA-CV-A6K1-01 | R |
| TCGA-CV-A6K2-01 | R |
| TCGA-CX-7082-01 | M0 |
| TCGA-CX-7085-01 | M0 |
| TCGA-CX-7086-01 | M0 |
| TCGA-CX-7219-01 | M0 |
| TCGA-D6-6515-01 | R |
| TCGA-D6-6516-01 | M0 |
| TCGA-D6-6517-01 | M0 |
| TCGA-D6-6823-01 | M0 |
| TCGA-D6-6824-01 | M0 |
| TCGA-D6-6825-01 | M0 |
| TCGA-D6-6826-01 | M0 |
| TCGA-D6-6827-01 | M0 |
| TCGA-D6-8568-01 | M0 |
| TCGA-D6-8569-01 | M0 |
| TCGA-D6-A4Z9-01 | M0 |
| TCGA-D6-A4ZB-01 | M0 |
| TCGA-D6-A6EK-01 | M0 |
| TCGA-D6-A6EM-01 | M0 |
| TCGA-D6-A6EO-01 | M0 |
| TCGA-D6-A6EP-01 | M0 |
| TCGA-D6-A6EQ-01 | M0 |
| TCGA-D6-A6ES-01 | M0 |
| TCGA-D6-A74Q-01 | M0 |
| TCGA-DQ-5624-01 | M0 |
| TCGA-DQ-5625-01 | R |
| TCGA-DQ-5629-01 | M0 |
| TCGA-DQ-5630-01 | M0 |
| TCGA-DQ-5631-01 | R |
| TCGA-DQ-7588-01 | R |
| TCGA-DQ-7589-01 | M1 |
| TCGA-DQ-7590-01 | M0 |
| TCGA-DQ-7591-01 | M0 |
| TCGA-DQ-7592-01 | M0 |
| TCGA-DQ-7593-01 | M0 |
| TCGA-DQ-7594-01 | M0 |
| TCGA-DQ-7595-01 | M0 |
| TCGA-DQ-7596-01 | R |
| TCGA-F7-7848-01 | M0 |
| TCGA-F7-8489-01 | M0 |
| TCGA-F7-A50G-01 | M0 |
| TCGA-F7-A50I-01 | M0 |
| TCGA-F7-A50J-01 | M0 |
| TCGA-F7-A61S-01 | R |
| TCGA-H7-7774-01 | R |
| TCGA-H7-8501-01 | M0 |
| TCGA-H7-A6C4-01 | M0 |
| TCGA-HD-7229-01 | M0 |
| TCGA-HD-7753-01 | M0 |
| TCGA-HD-7754-01 | M0 |
| TCGA-HD-7831-01 | M0 |
| TCGA-HD-7832-01 | M0 |
| TCGA-HD-7917-01 | M0 |
| TCGA-HD-8224-01 | R |
| TCGA-HD-8314-01 | M0 |
| TCGA-HD-A633-01 | R |
| TCGA-HL-7533-01 | M0 |
| TCGA-IQ-7630-01 | M0 |
| TCGA-IQ-7631-01 | M0 |
| TCGA-IQ-7632-01 | M0 |
| TCGA-IQ-A61I-01 | M0 |
| TCGA-IQ-A61J-01 | M0 |
| TCGA-KU-A66S-01 | R |
| TCGA-KU-A66T-01 | R |
| TCGA-KU-A6H7-01 | M0 |
| TCGA-KU-A6H8-01 | R |
| TCGA-MT-A51W-01 | M0 |
| TCGA-MT-A51X-01 | M0 |
| TCGA-MT-A67D-01 | M0 |
| TCGA-MT-A7BN-01 | R |
| TCGA-MZ-A5BI-01 | M0 |
| TCGA-MZ-A6I9-01 | R |
| TCGA-P3-A5Q6-01 | R |
| TCGA-P3-A6T5-01 | R |
| TCGA-P3-A6T6-01 | R |
| TCGA-QK-A64Z-01 | R |
| TCGA-QK-A6IF-01 | R |
| TCGA-QK-A6IG-01 | R |
| TCGA-QK-A6IH-01 | R |
| TCGA-QK-A6II-01 | R |
| TCGA-QK-A6IJ-01 | M0 |
| TCGA-QK-A6V9-01 | M0 |
| TCGA-QK-A6VB-01 | M0 |
| TCGA-QK-A6VC-01 | M0 |
| TCGA-RS-A6TO-01 | R |
| TCGA-RS-A6TP-01 | M0 |
| TCGA-T2-A6WX-01 | M0 |
| TCGA-T2-A6WZ-01 | R |
| TCGA-T2-A6X0-01 | M0 |
| TCGA-T2-A6X2-01 | M0 |
| TCGA-TN-A7HI-01 | M0 |
| TCGA-TN-A7HJ-01 | M0 |
| TCGA-TN-A7HL-01 | M0 |
| TCGA-UF-A718-01 | M0 |
| TCGA-UF-A719-01 | M0 |
| TCGA-UF-A71A-01 | M0 |
| TCGA-UF-A71B-01 | M0 |
| TCGA-UF-A71D-01 | M0 |
| TCGA-UF-A71E-01 | M0 |
| TCGA-UF-A7J9-01 | M0 |
| TCGA-UF-A7JA-01 | M0 |
| TCGA-UF-A7JC-01 | M0 |
| TCGA-UF-A7JD-01 | M0 |
| TCGA-UF-A7JF-01 | M0 |
| TCGA-UF-A7JH-01 | M0 |
| TCGA-UF-A7JJ-01 | M0 |
| TCGA-UF-A7JK-01 | M0 |
| TCGA-UF-A7JO-01 | M0 |
| TCGA-UF-A7JS-01 | R |
| TCGA-UF-A7JT-01 | M0 |
| TCGA-UF-A7JV-01 | M0 |
| TCGA-WA-A7GZ-01 | M0 |
| TCGA-WA-A7H4-01 | M0 |
| TCGA-CV-5430-11 | N |
| TCGA-CV-5431-11 | N |
| TCGA-CV-5432-11 | N |
| TCGA-CV-5434-11 | N |
| TCGA-CV-5435-11 | N |
| TCGA-CV-5436-11 | N |
| TCGA-CV-5439-11 | N |
| TCGA-CV-5440-11 | N |
| TCGA-CV-5441-11 | N |
| TCGA-CV-5442-11 | N |
| TCGA-CV-5443-11 | N |
| TCGA-CV-5444-11 | N |
| TCGA-CV-5966-11 | N |
| TCGA-CV-5970-11 | N |
| TCGA-CV-5971-11 | N |
| TCGA-CV-5973-11 | N |
| TCGA-CV-5976-11 | N |
| TCGA-CV-5977-11 | N |
| TCGA-CV-5978-11 | N |
| TCGA-CV-5979-11 | N |
| TCGA-CV-6003-11 | N |
| TCGA-CV-6433-11 | N |
| TCGA-CV-6436-11 | N |
| TCGA-CV-6441-11 | N |
| TCGA-CV-6933-11 | N |
| TCGA-CV-6934-11 | N |
| TCGA-CV-6935-11 | N |
| TCGA-CV-6936-11 | N |
| TCGA-CV-6938-11 | N |
| TCGA-CV-6939-11 | N |
| TCGA-CV-6943-11 | N |
| TCGA-CV-6951-11 | N |
| TCGA-CV-6952-11 | N |
| TCGA-CV-6953-11 | N |
| TCGA-CV-6954-11 | N |
| TCGA-CV-6955-11 | N |
| TCGA-CV-6956-11 | N |
| TCGA-CV-6959-11 | N |
| TCGA-CV-6960-11 | N |
| TCGA-CV-6961-11 | N |
| TCGA-CV-6962-11 | N |
| TCGA-CV-7089-11 | N |
| TCGA-CV-7091-11 | N |
| TCGA-CV-7097-11 | N |
| TCGA-CV-7101-11 | N |
| TCGA-CV-7103-11 | N |
| TCGA-CV-7177-11 | N |
| TCGA-CV-7178-11 | N |
| TCGA-CV-7183-11 | N |
| TCGA-CV-7235-11 | N |
| TCGA-CV-7238-11 | N |
| TCGA-CV-7242-11 | N |
| TCGA-CV-7245-11 | N |
| TCGA-CV-7250-11 | N |
| TCGA-CV-7252-11 | N |
| TCGA-CV-7255-11 | N |
| TCGA-CV-7261-11 | N |
| TCGA-CV-7263-11 | N |
| TCGA-CV-7406-11 | N |
| TCGA-CV-7416-11 | N |
| TCGA-CV-7423-11 | N |
| TCGA-CV-7424-11 | N |
| TCGA-CV-7425-11 | N |
| TCGA-CV-7432-11 | N |
| TCGA-CV-7434-11 | N |
| TCGA-CV-7437-11 | N |
| TCGA-CV-7438-11 | N |
| TCGA-CV-7440-11 | N |
| TCGA-H7-A6C5-11 | N |
| TCGA-HD-8635-11 | N |
| TCGA-HD-A6HZ-11 | N |
| TCGA-HD-A6I0-11 | N |
| TCGA-WA-A7GZ-11 | N |

**Supplementary Table S2:**
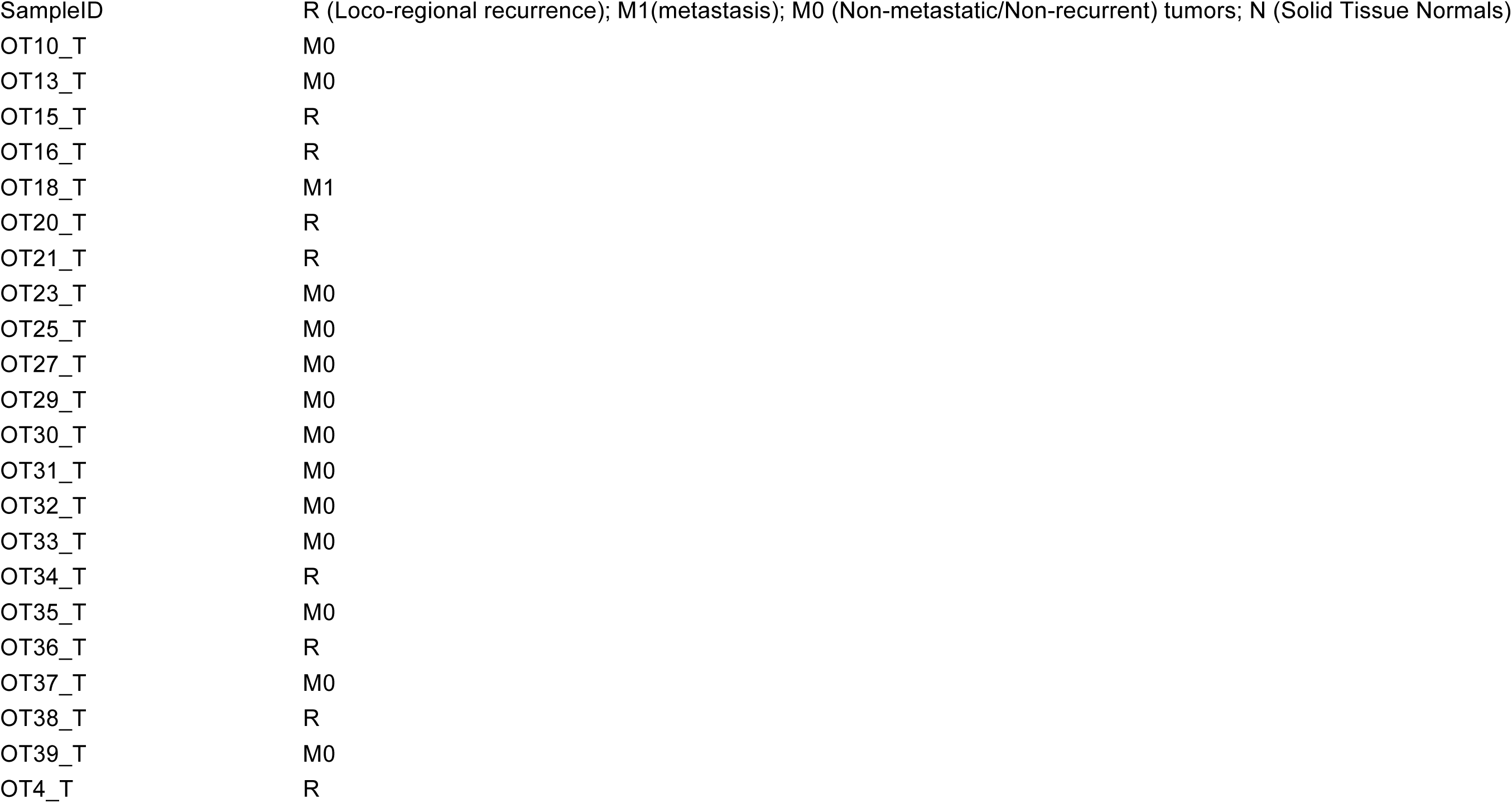
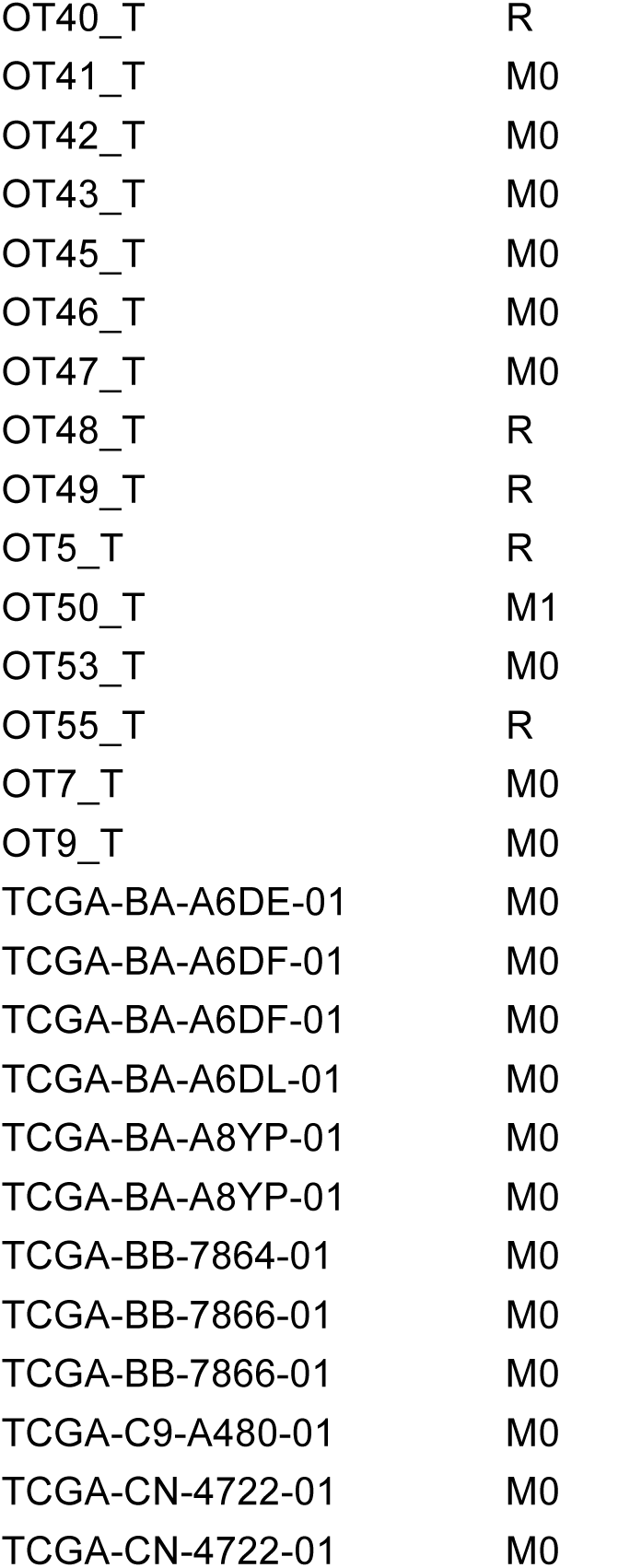
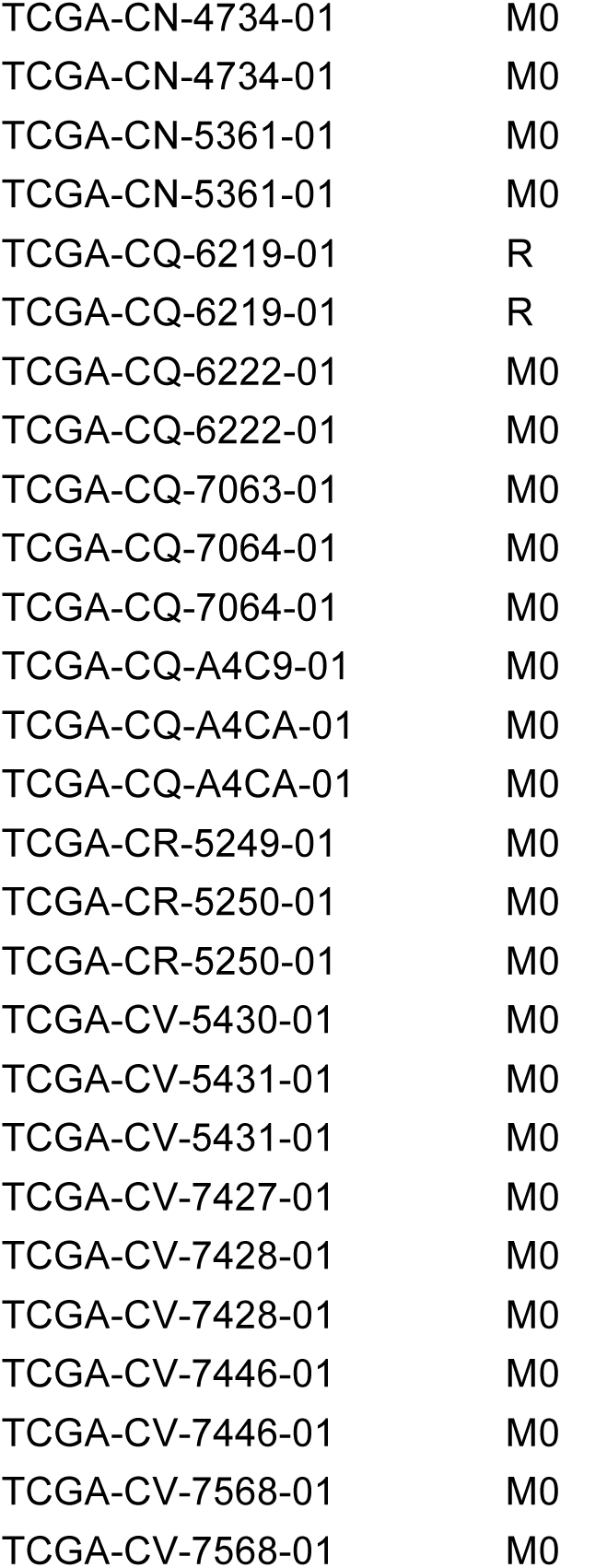
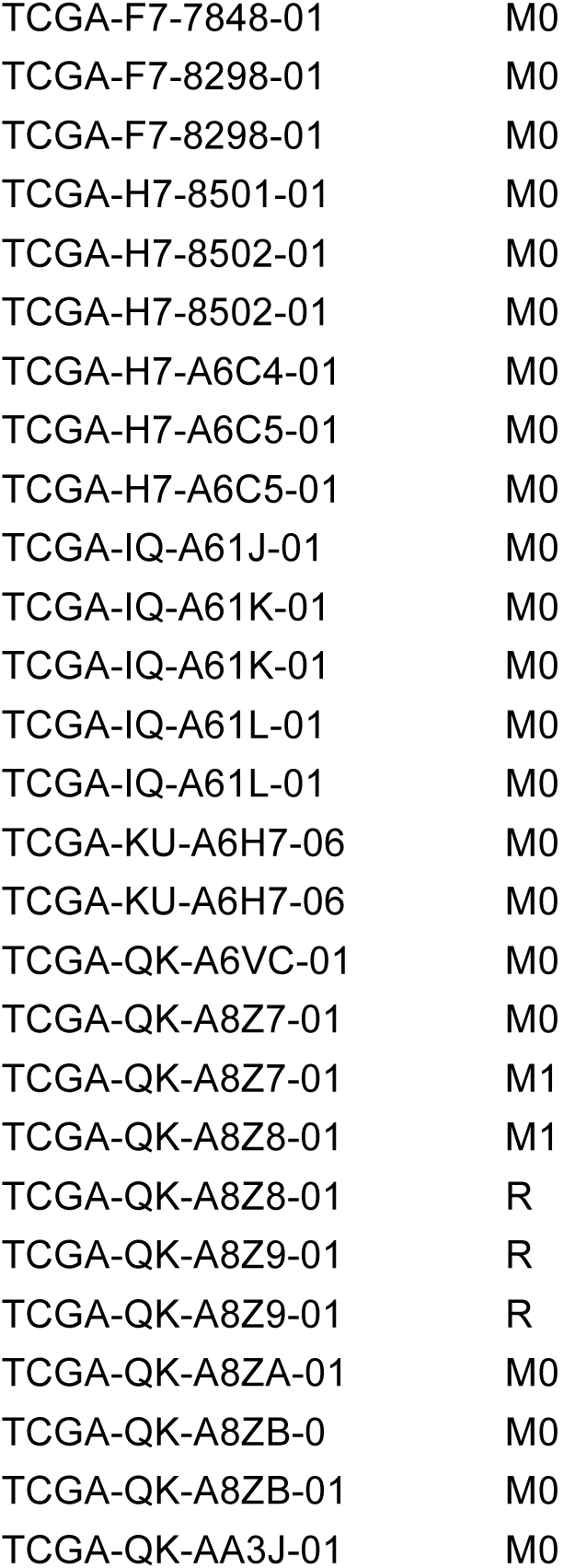
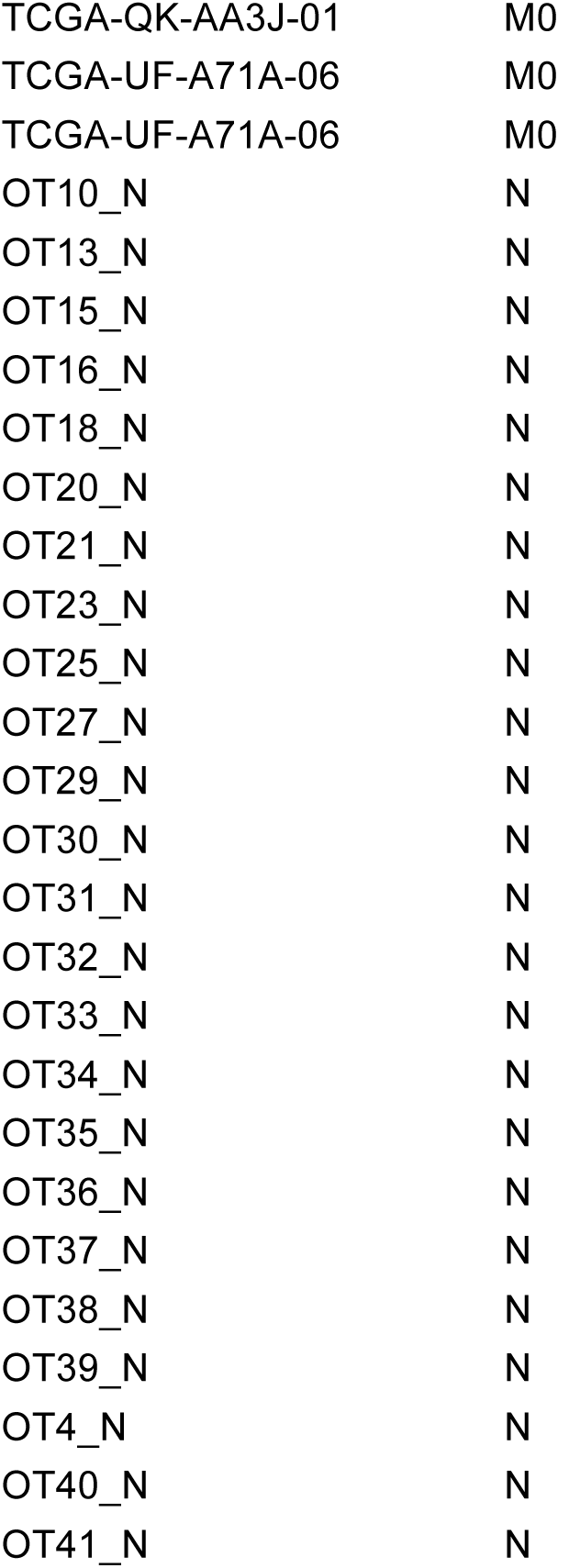
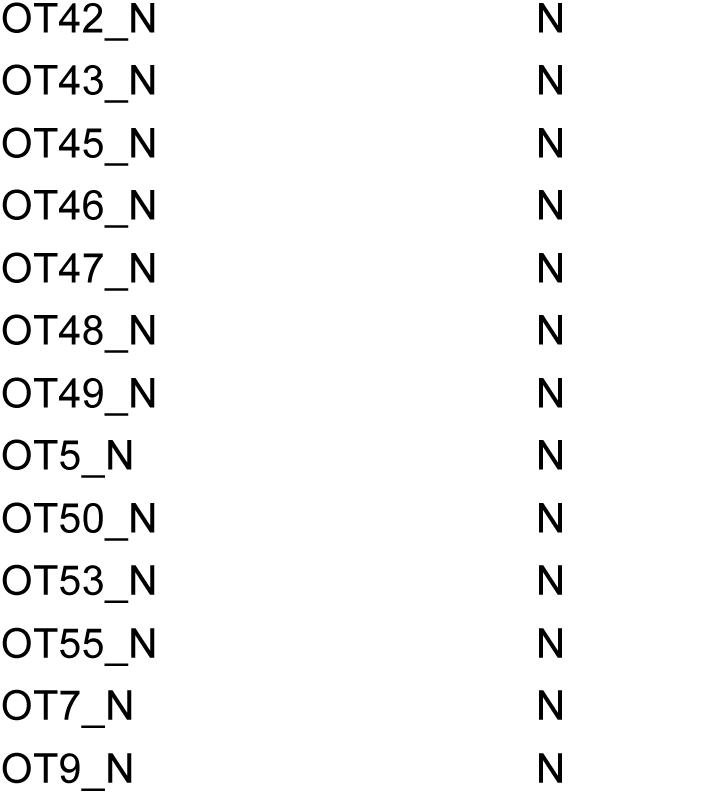
Oral tongue squamous cell carcinoma (OTSCC) and TCGA HNSCC samples lacking cross-platform overlap, and their clinical attributes used for confirmation of MR44 discovery.

**Supplementary Table S3:** Oral tongue squamous cell carcinoma (OTSCC) (*n* = 18) with all four events assayed within the same tumor, and their clinical attributes used for the validation.

| Supplementary Table S3: Oral tongue squamous cell carcinoma (OTSCC) with all four events assayed within the same tumor, and their clinical attributes used for validation. |  |
| --- | --- |
| S<br>a<br>m<br>p<br>l<br>e<br>I<br>D | R (Loco-regional recurrence); M1(metastasis); M0 (Non-metastatic/Non-recurrent) tumors; N (Solid Tissue Normals). |
| OT1_T | M0 |
| OT11_T | M0 |
| OT12_T | M0 |
| OT14_T | M0 |
| OT17_T | M0 |
| OT19_T | M0 |
| OT2_T | M1 |
| OT22_T | M0 |
| OT24_T | M0 |
| OT26_T | M0 |
| OT28_T | M0 |
| OT52_T | M0 |
| OT54_T | M0 |
| OT3_T | M0 |
| OT6_T | M0 |
| OT8_T | M0 |
| OT44_T | R |
| OT51_T | M1 |
| OT1_N | N |
| OT11_N | N |
| OT12_N | N |
| OT14_N | N |
| OT17_N | N |
| OT19_N | N |
| OT2_N | N |
| OT22_N | N |
| OT24_N | N |
| OT26_N | N |
| OT28_N | N |
| OT52_N | N |
| OT54_N | N |
| OT3_N | N |
| OT6_N | N |
| OT8_N | N |
| OT44_N | N |
| OT51_N | N |

**Supplementary Table S4:**
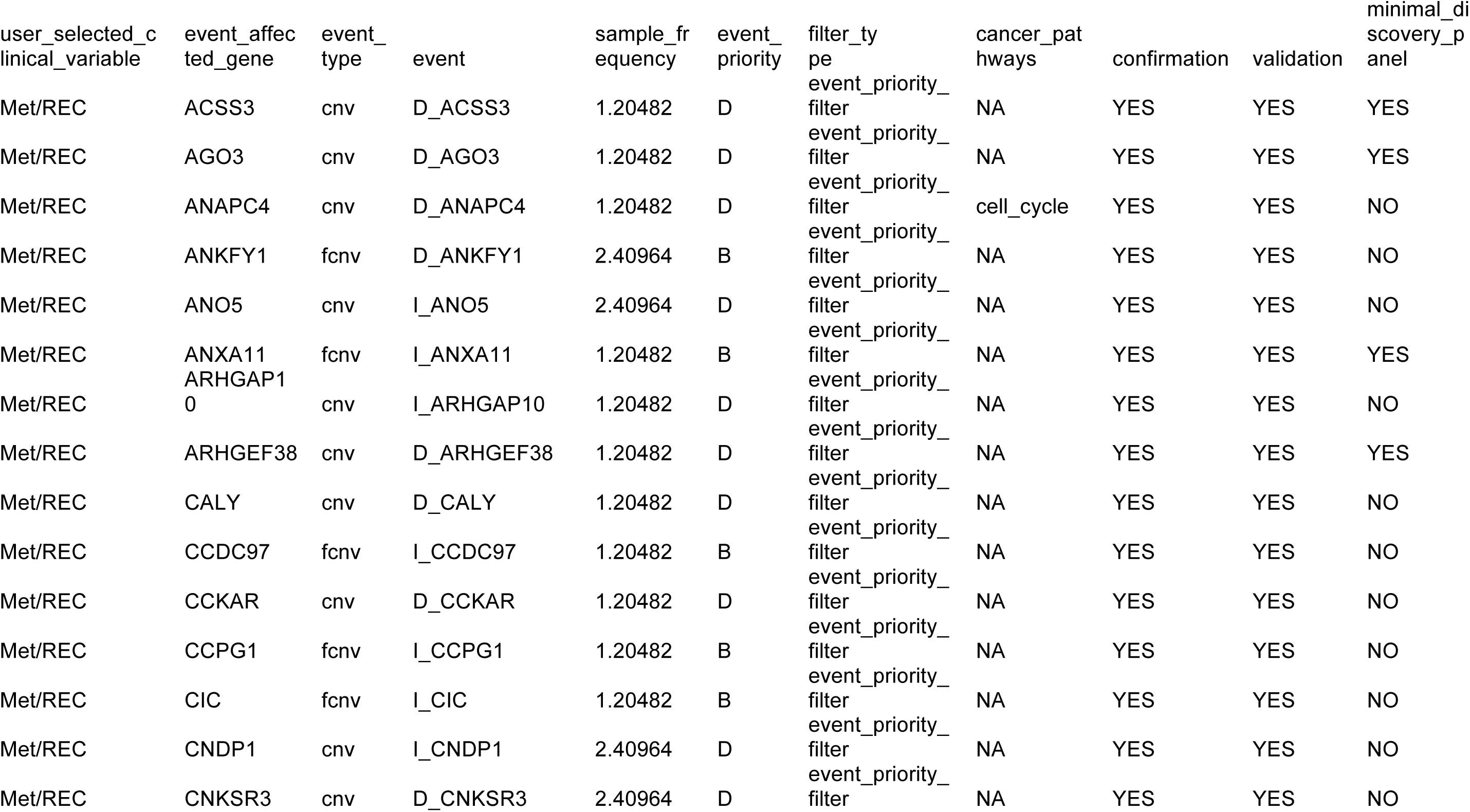
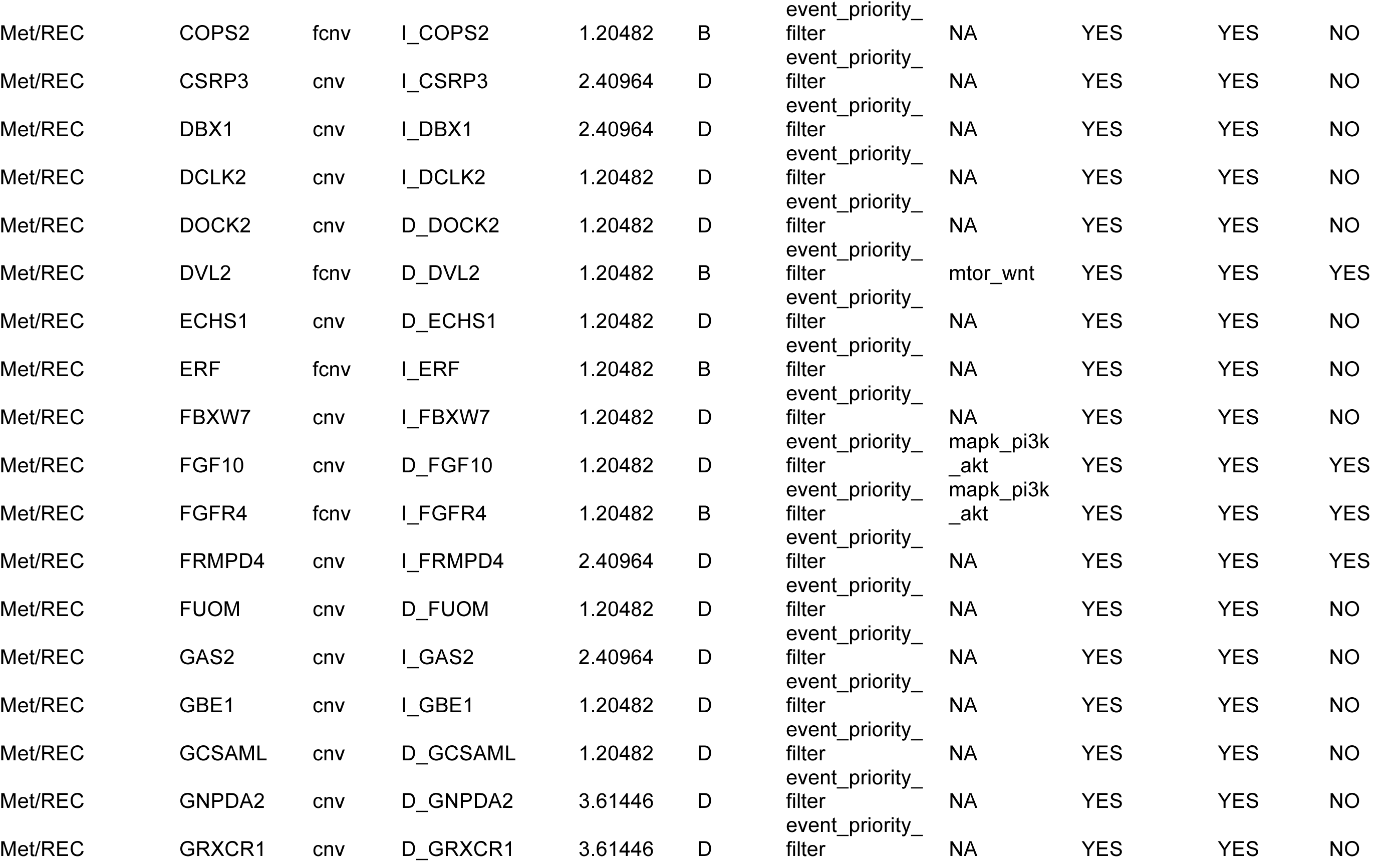
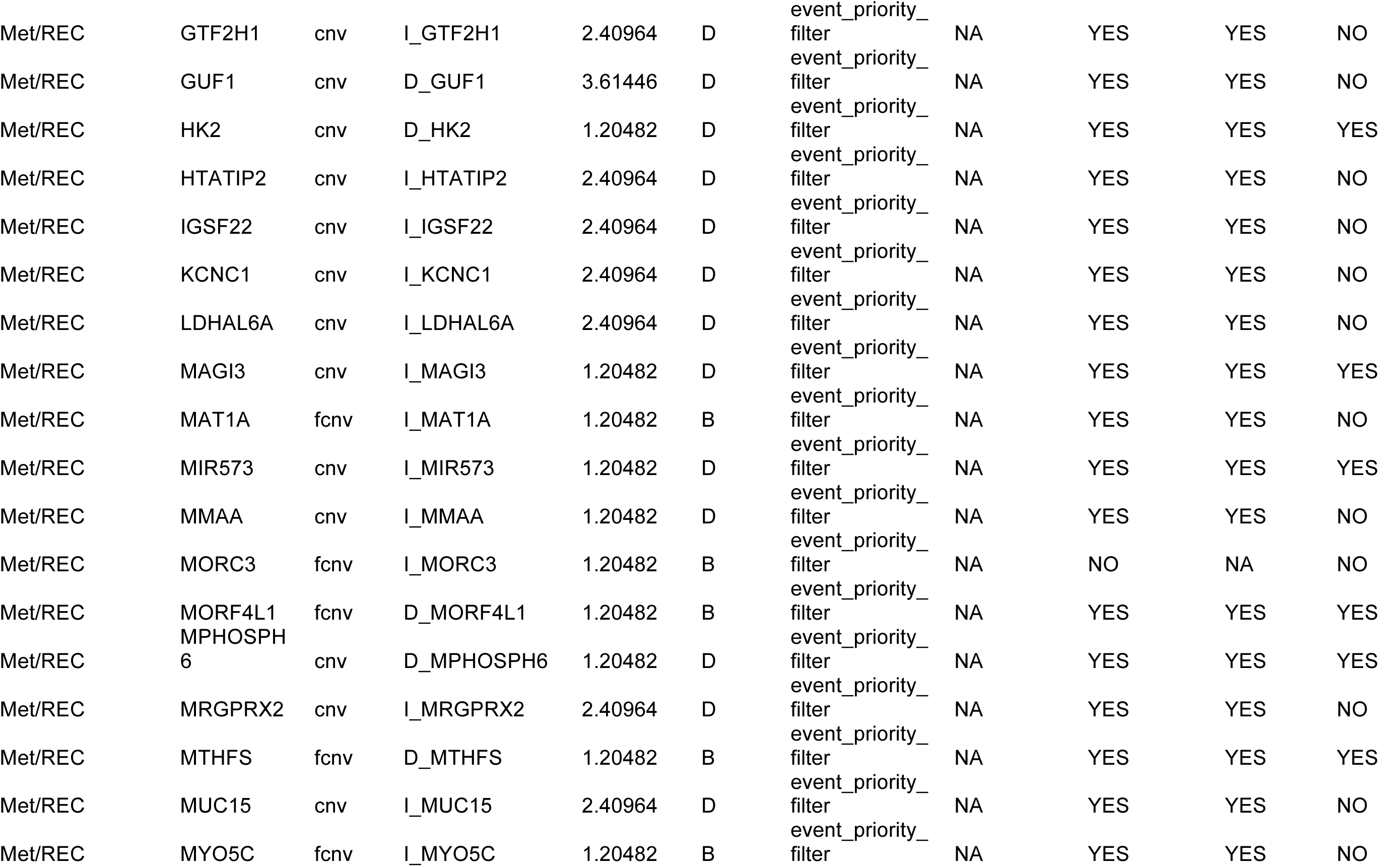
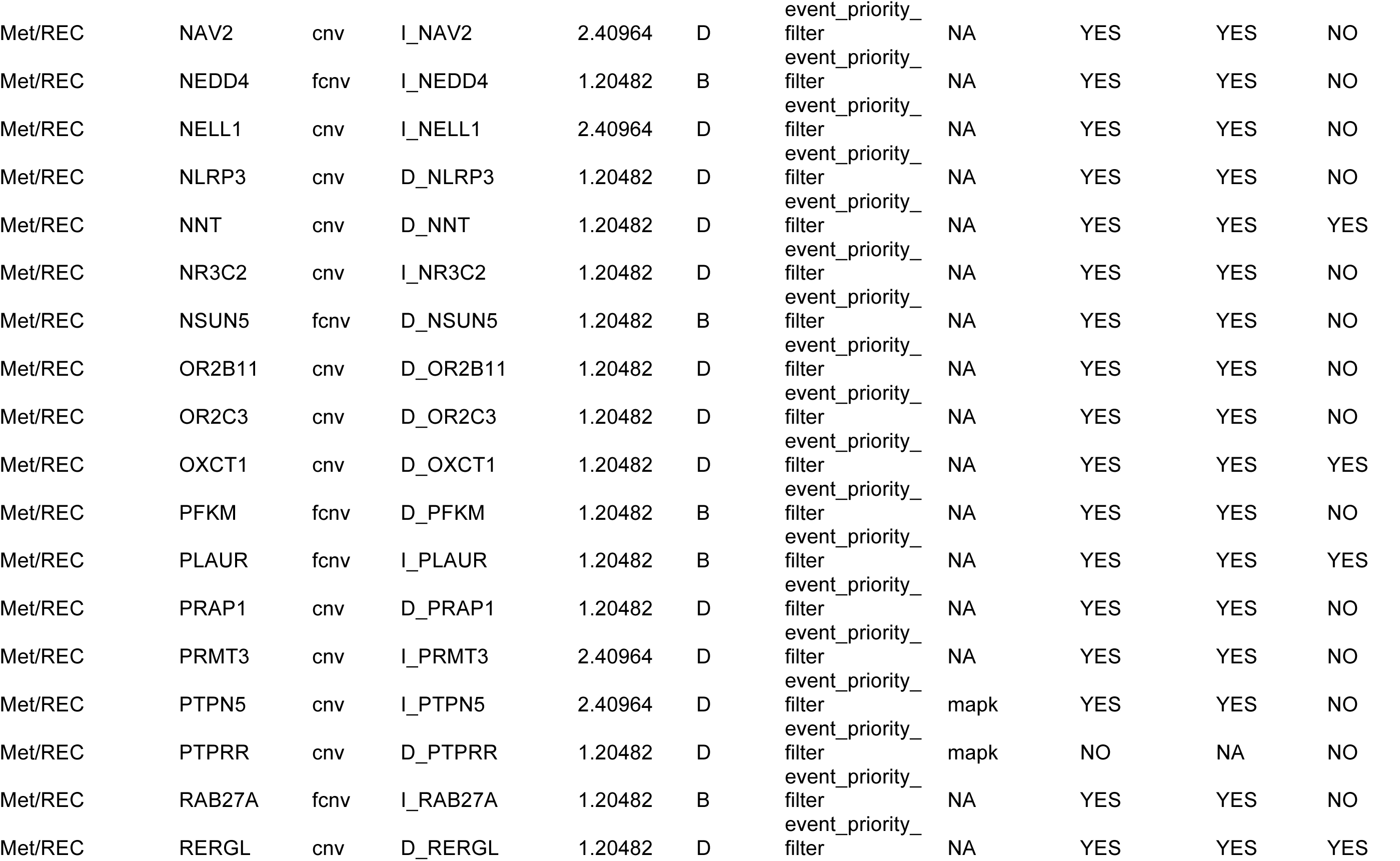
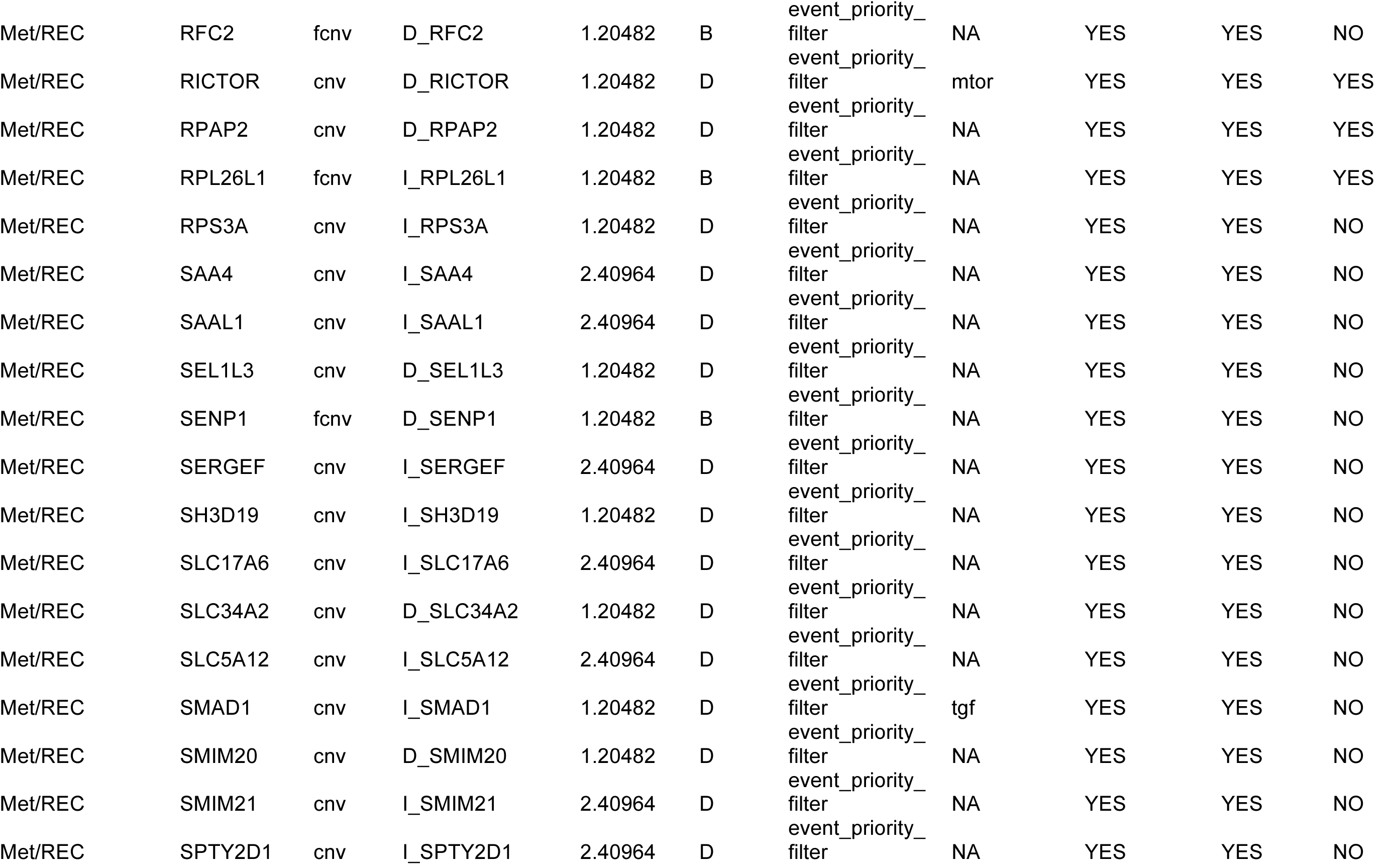
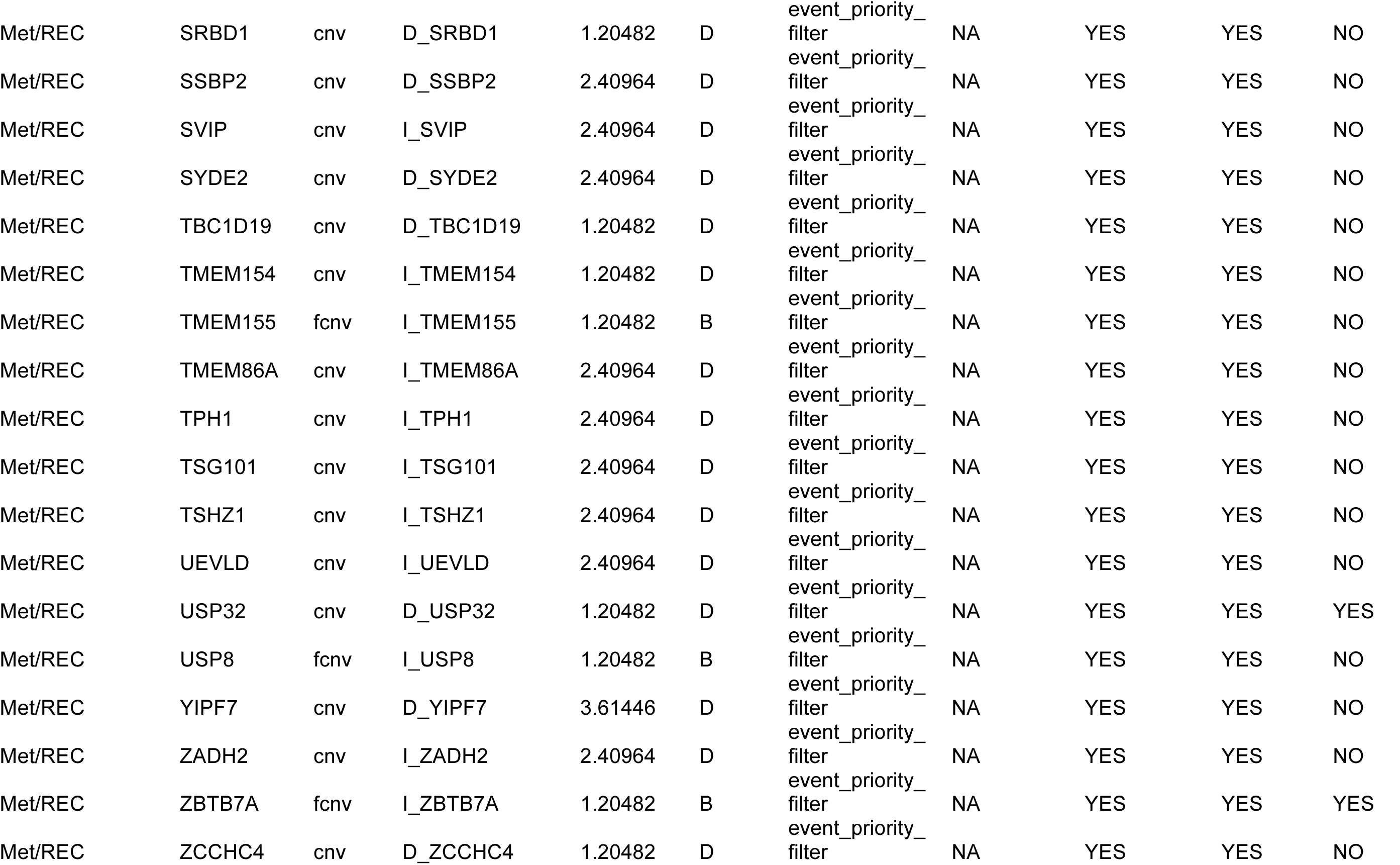
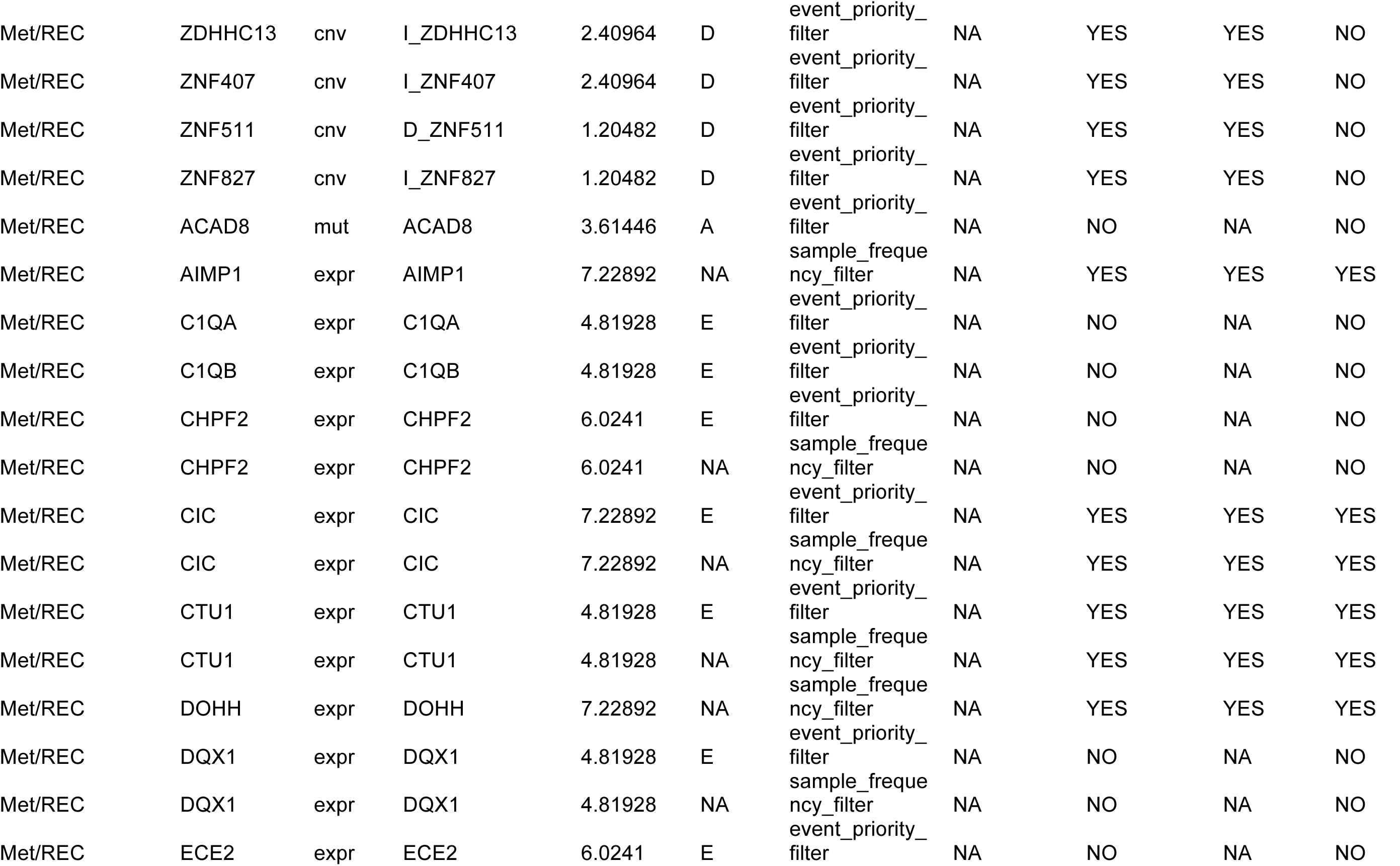
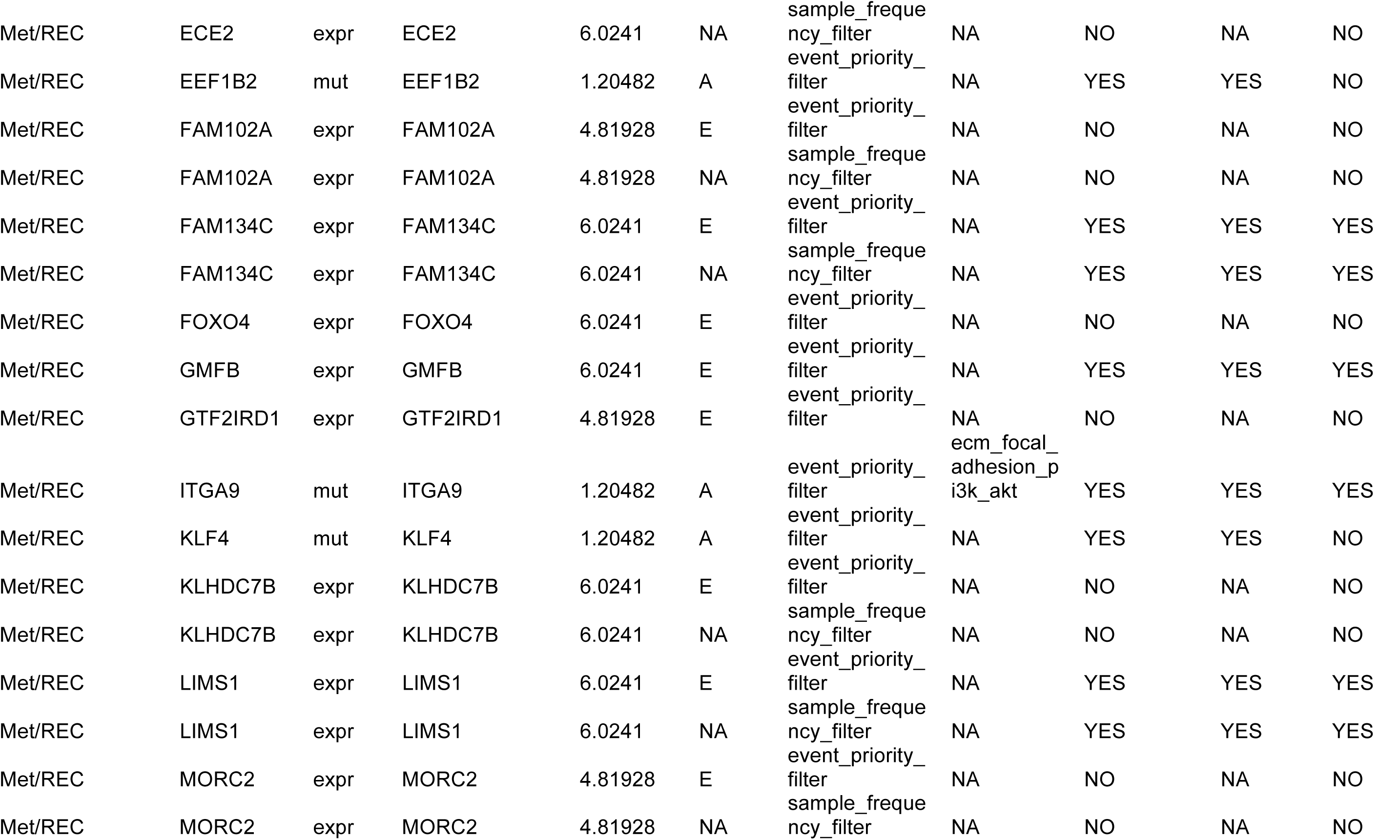
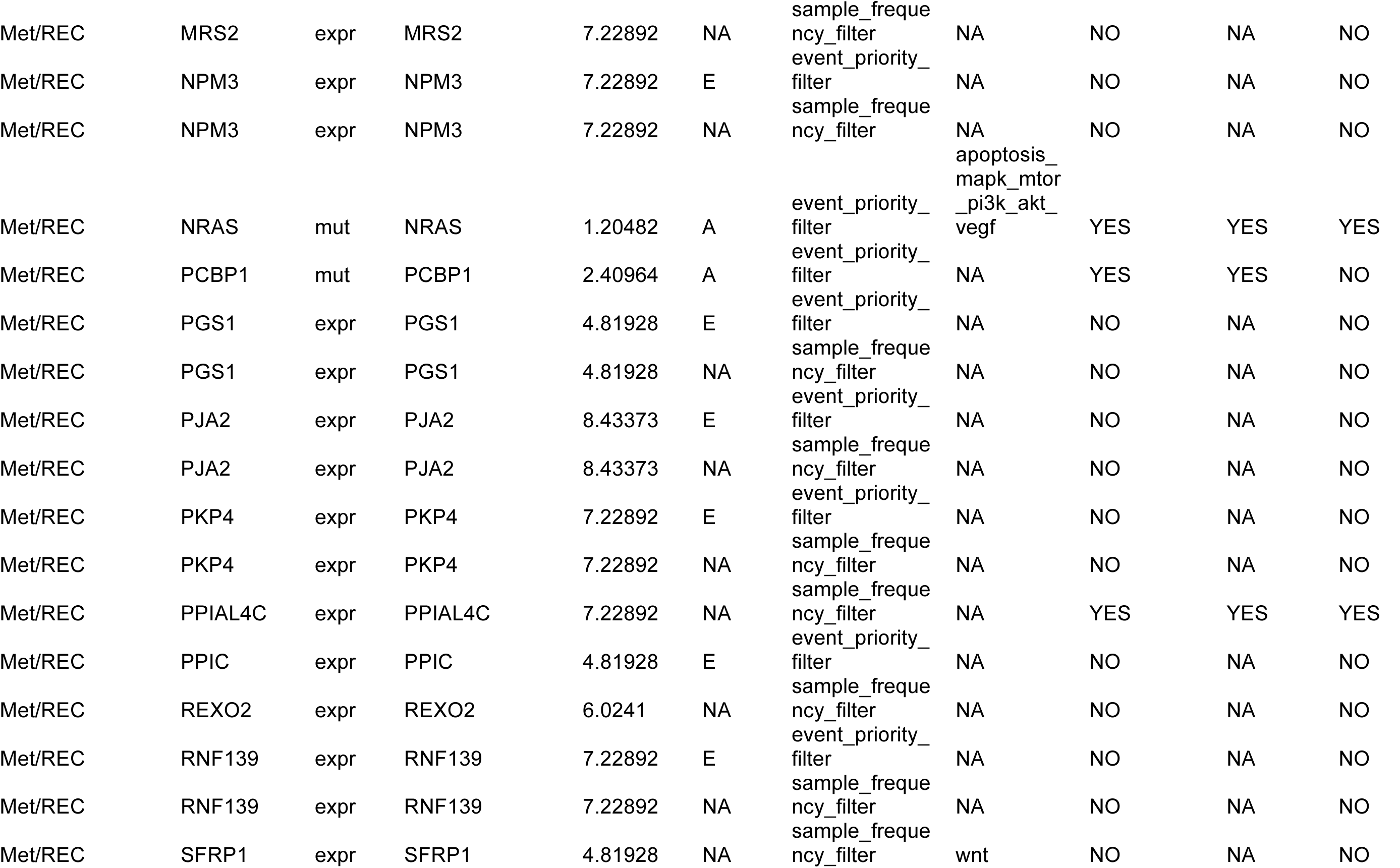
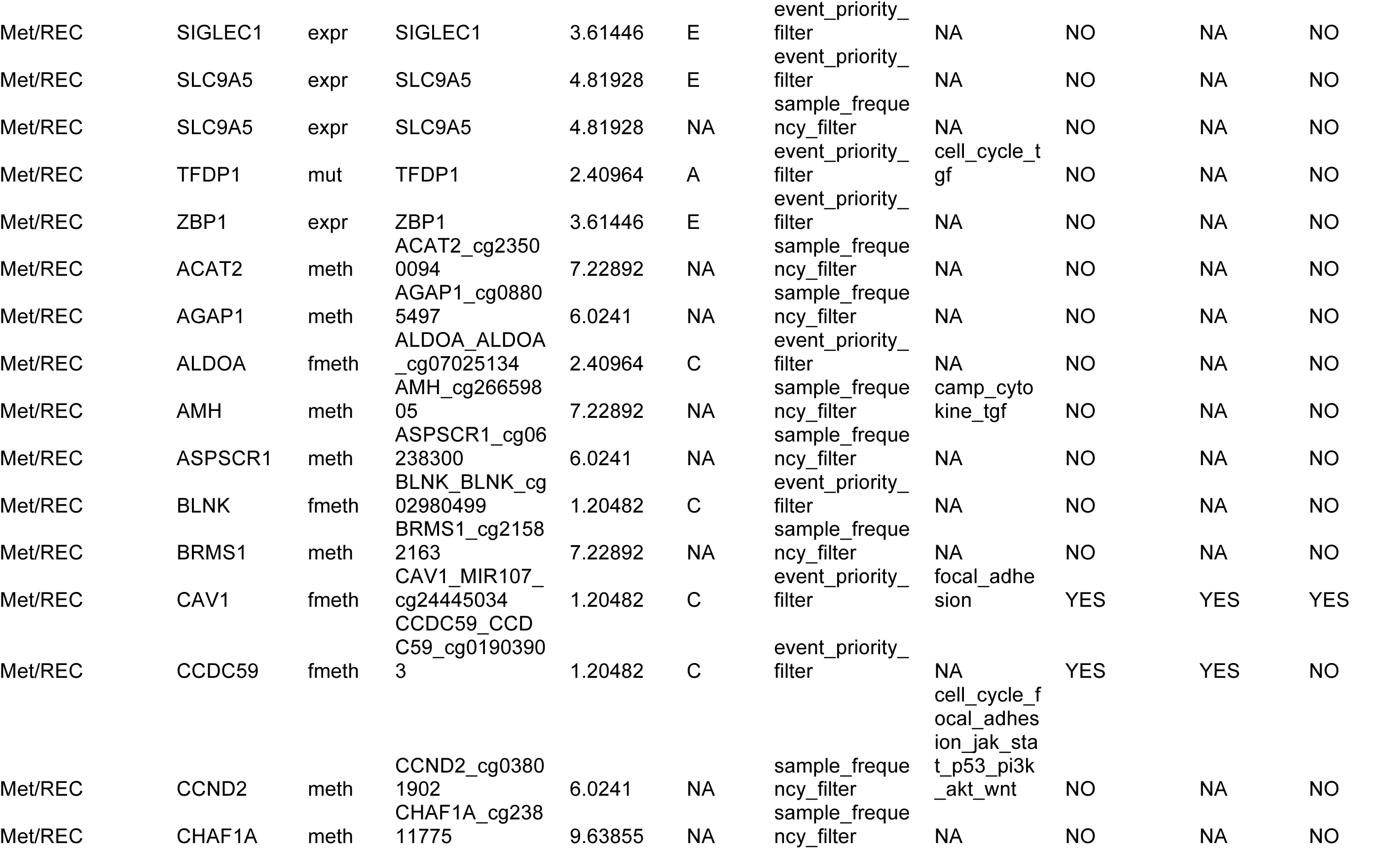
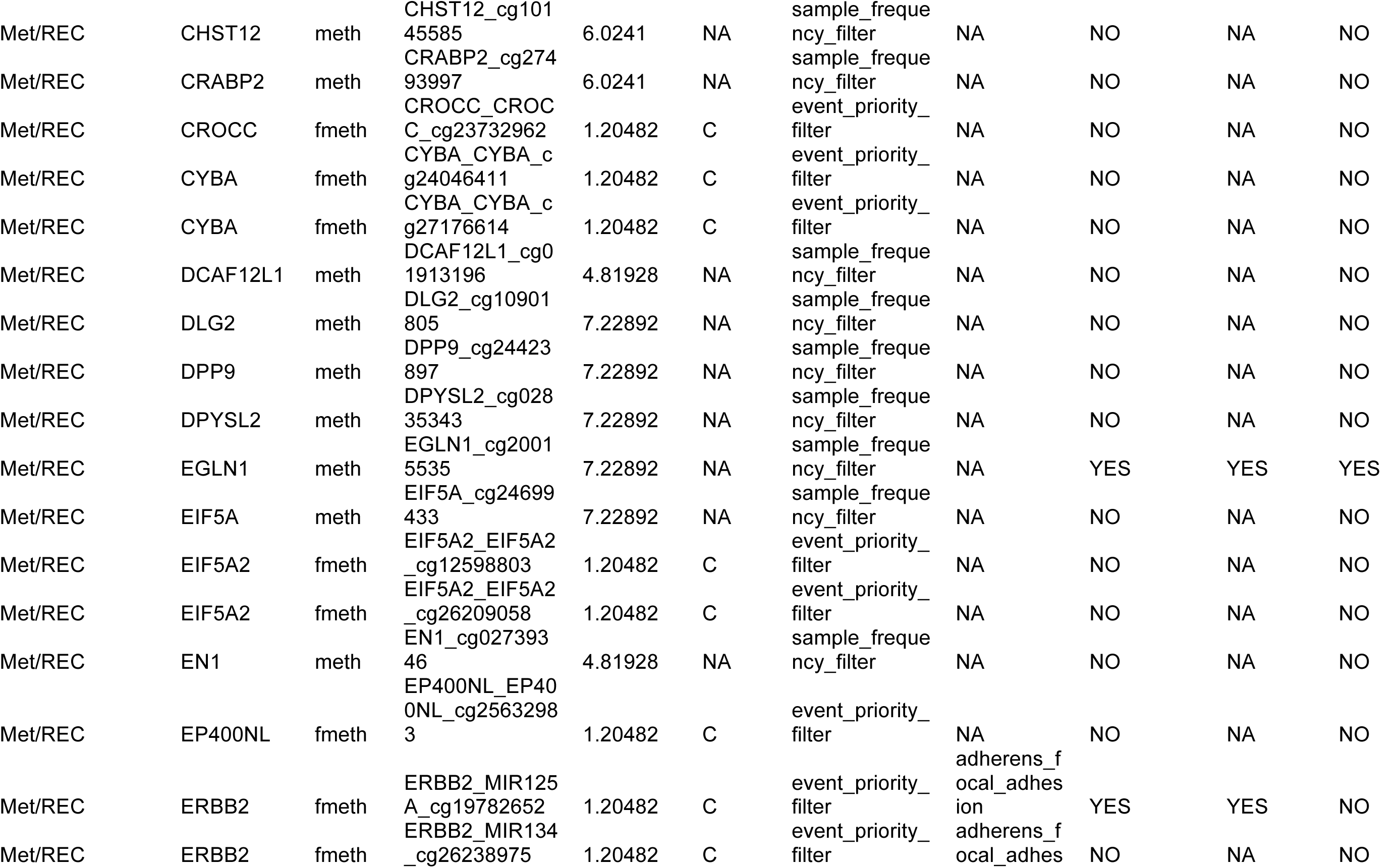
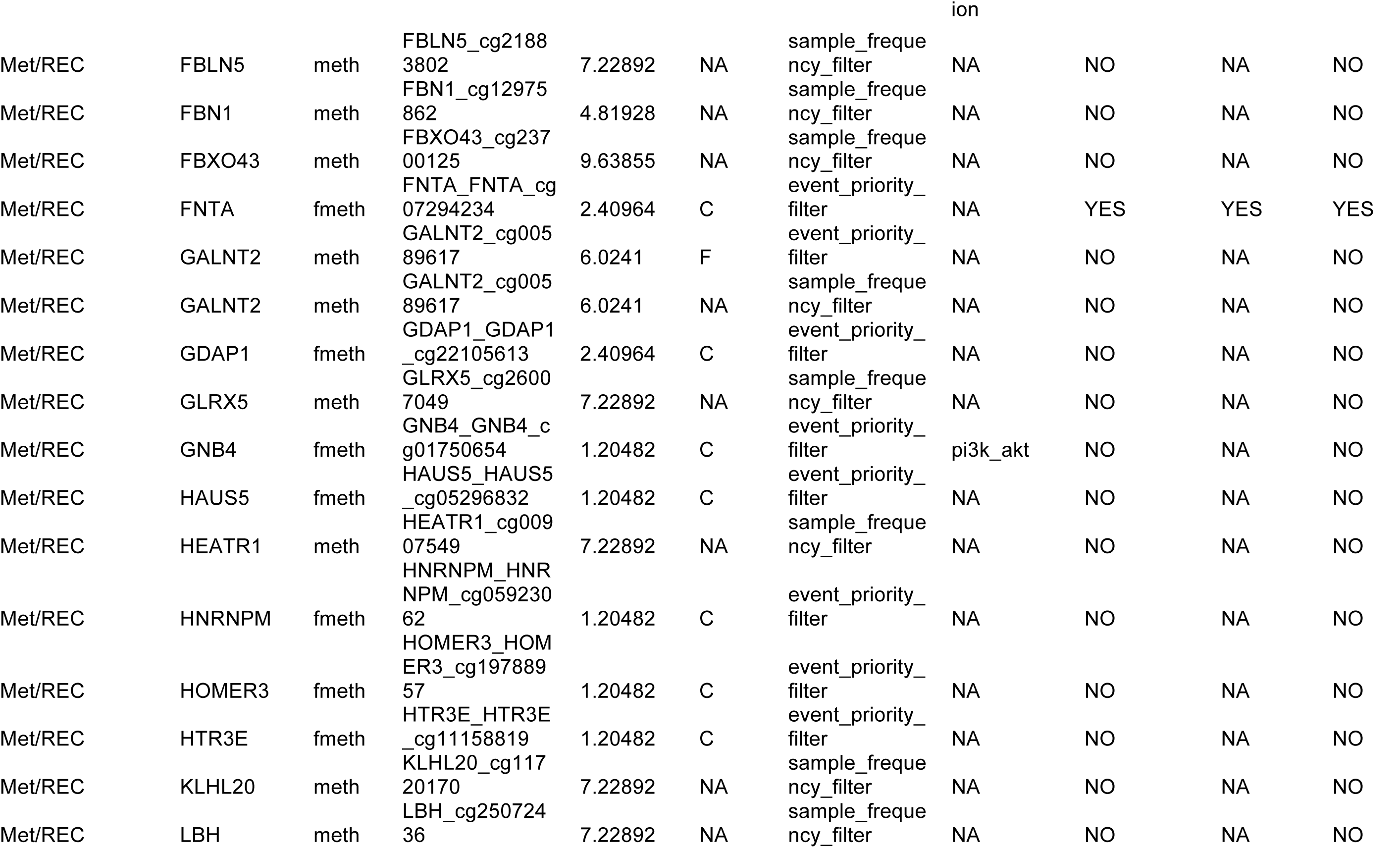
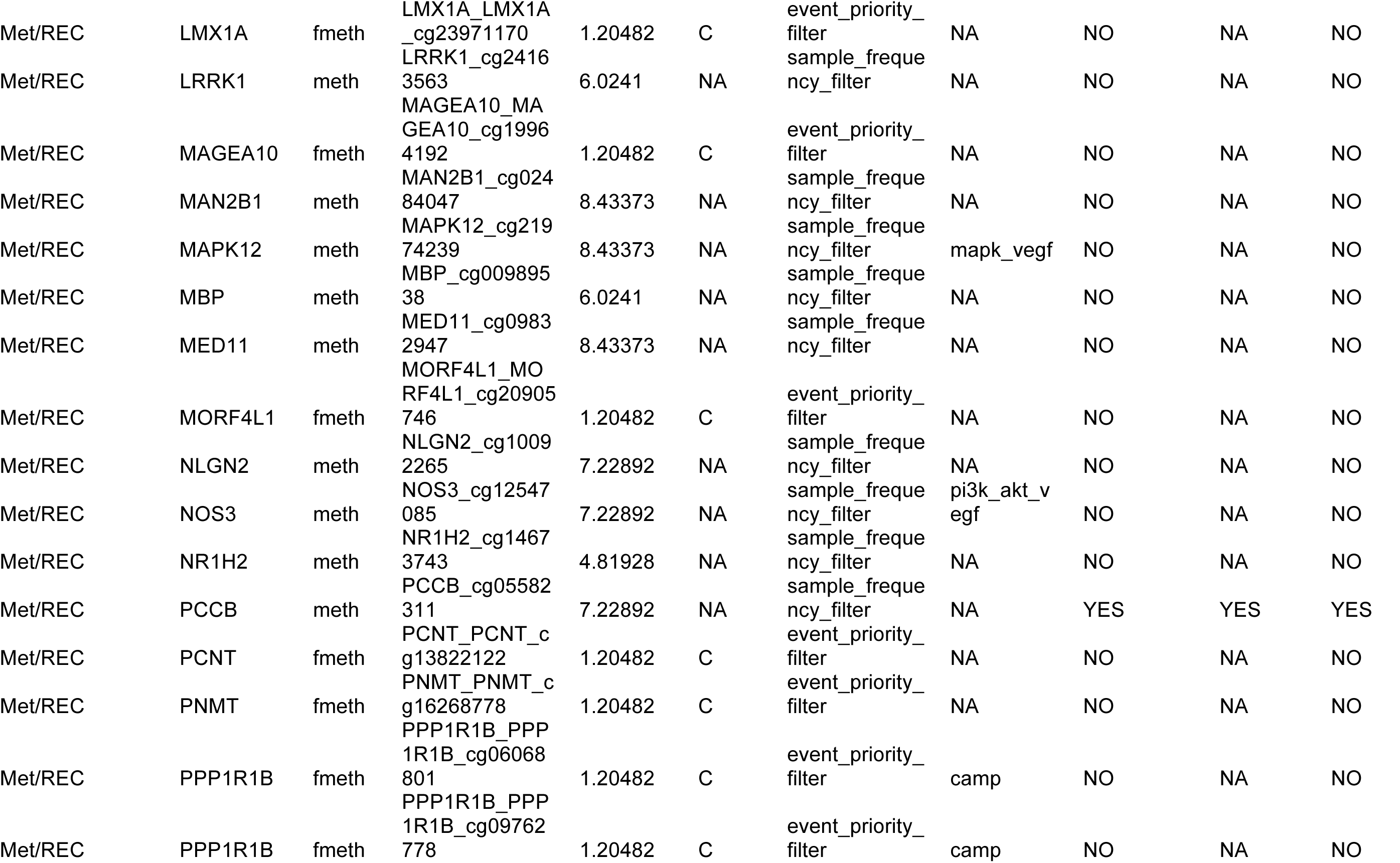
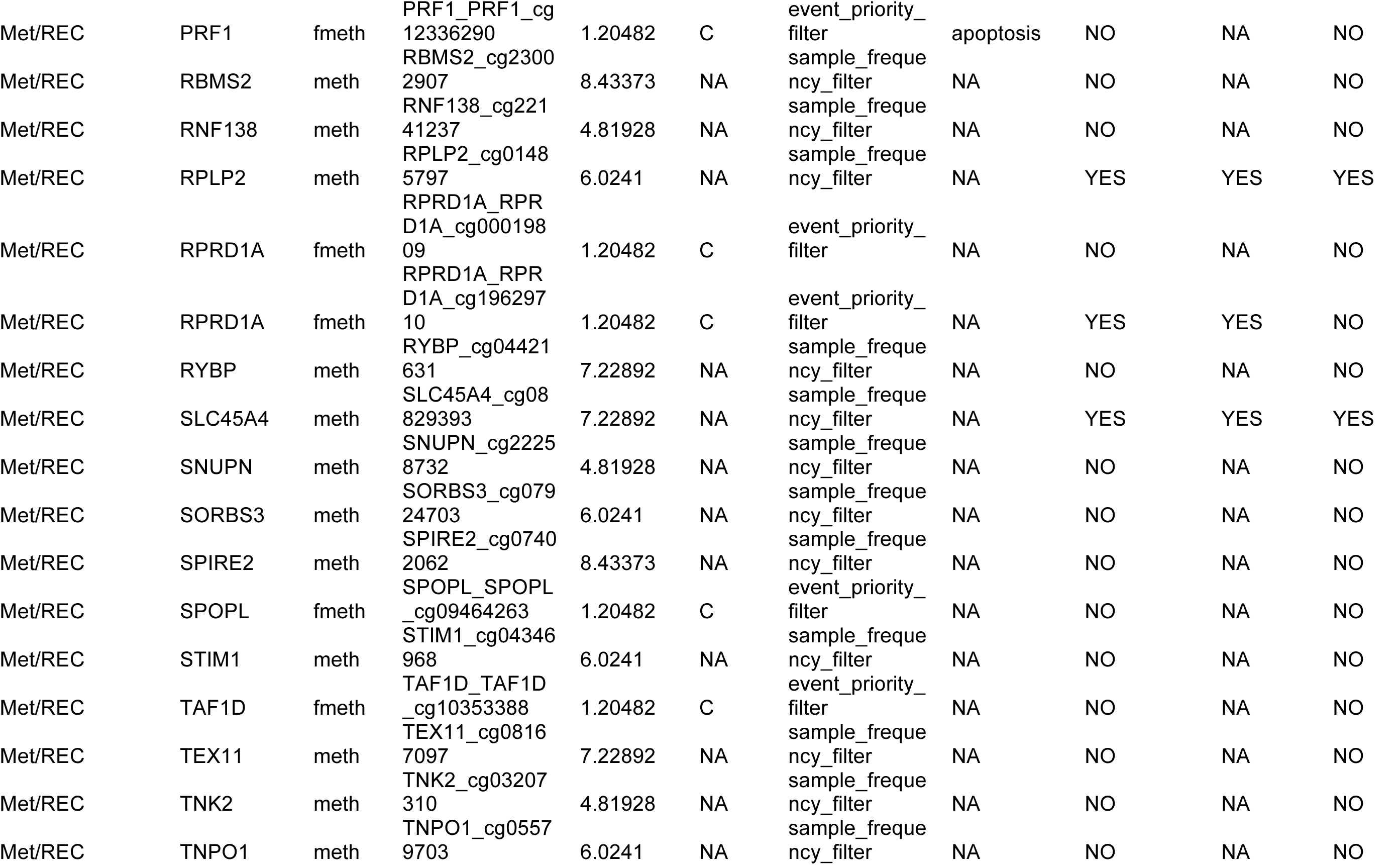
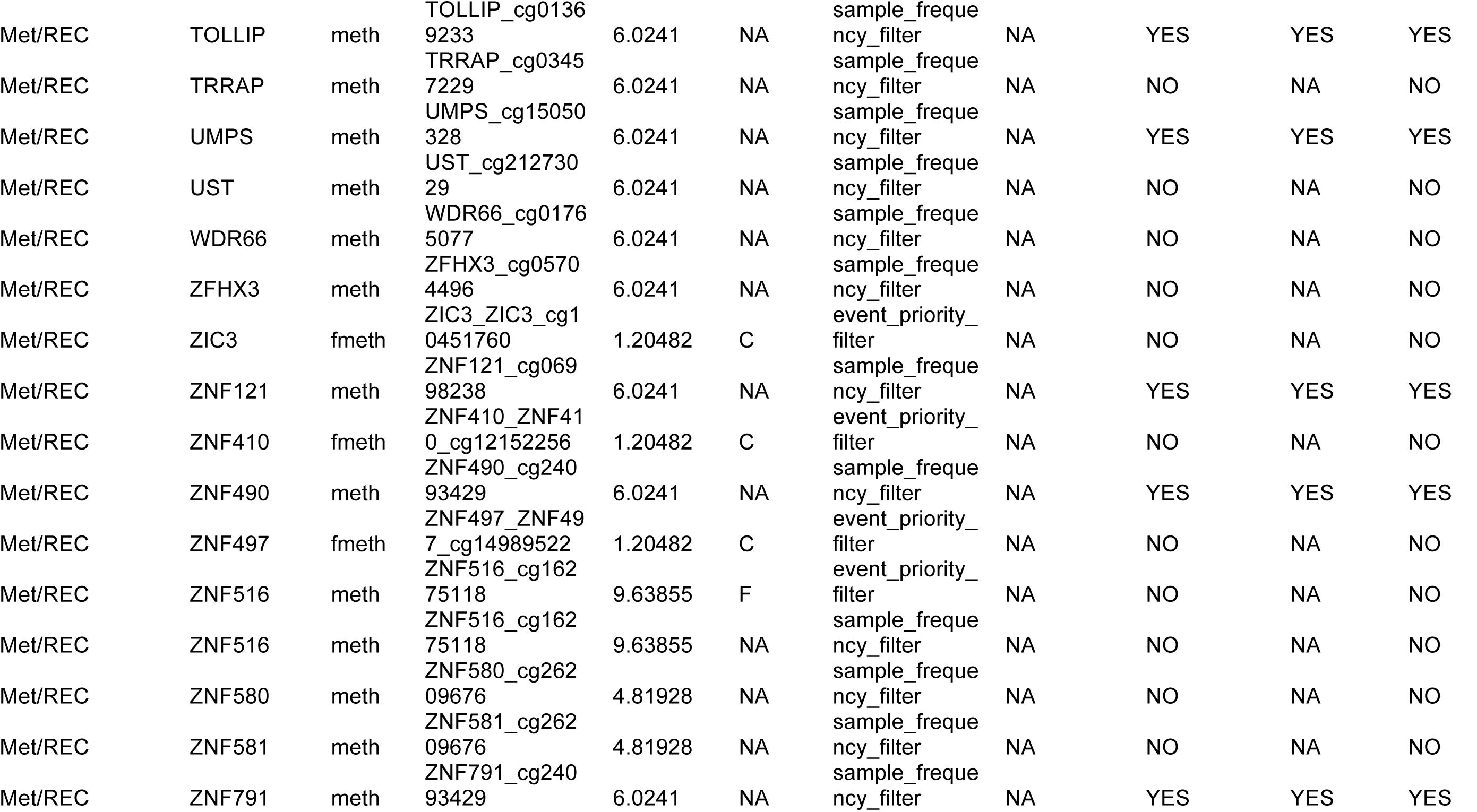
Discovered events associated with Metastases and Recurrence in HNSCC, pathways mapped, confirmation and validation status.

**Supplementary Figure S1:**
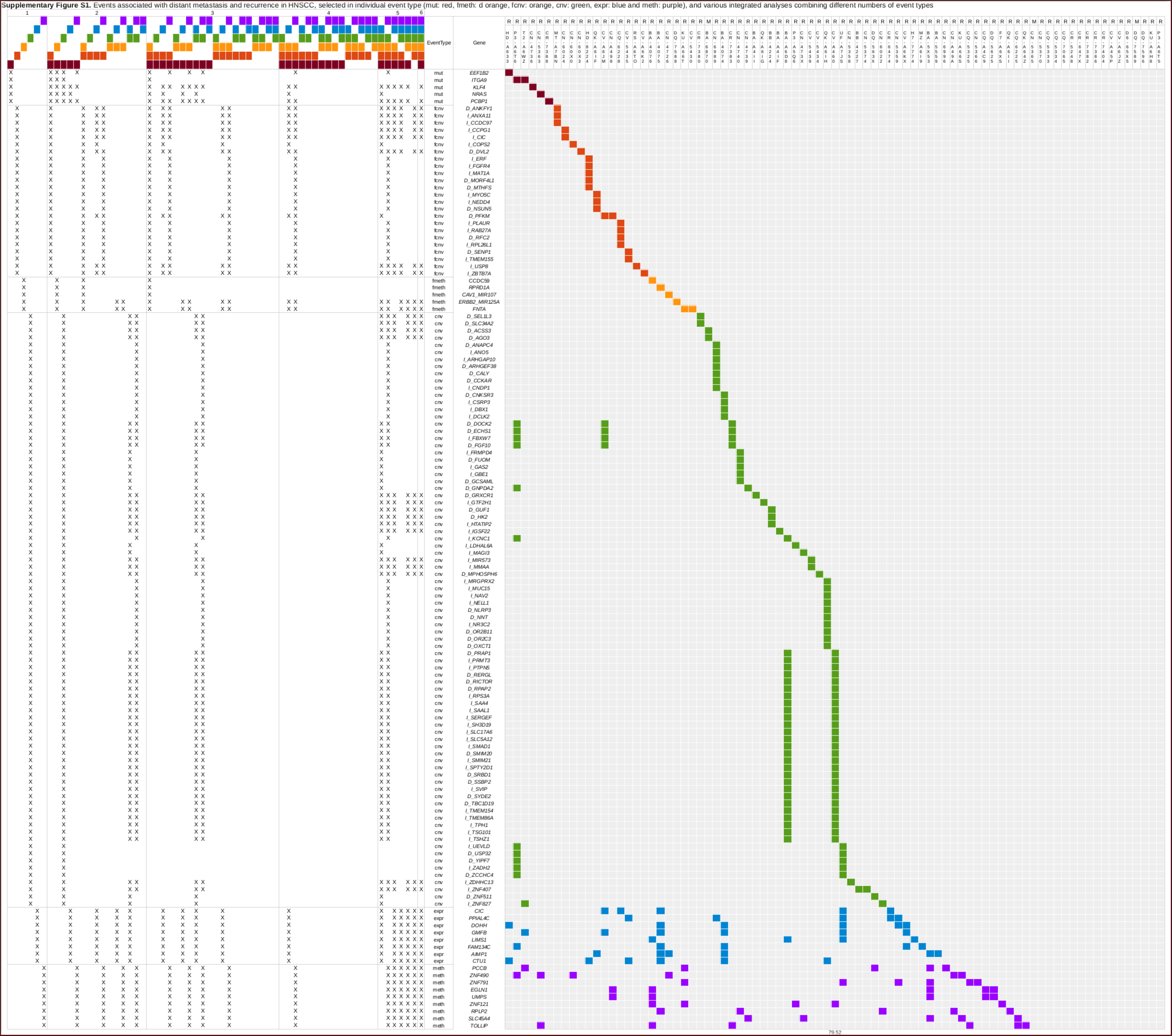
Events associated with distant metastasis and recurrence in HNSCC with single-event or multi-event signatures, selected in individual event type (mut: red, fmeth: d orange, fcnv: orange, cnv: green, expr: blue and meth: purple) and various integrated analyses, combining different numbers of event types.

